# Proteogenomic analysis unveils the HLA Class I presented immunopeptidome in melanoma and EGFR mutant lung adenocarcinoma

**DOI:** 10.1101/2020.08.04.236331

**Authors:** Yue A. Qi, Tapan K. Maity, Constance M. Cultraro, Vikram Misra, Xu Zhang, Catherine Ade, Shaojian Gao, David Milewski, Khoa D. Nguyen, Mohammad H. Ebrahimabadi, Ken-ichi Hanada, Javed Khan, Cenk Sahinalp, James C. Yang, Udayan Guha

## Abstract

Immune checkpoint inhibitor and adoptive lymphocyte transfer-based therapies have shown great therapeutic potential for cancers with high tumor mutation burden (TMB). Here, we employed mass spectrometry (MS)-based proteogenomic large-scale profiling to identify potential immunogenic human leukocyte antigen (HLA) Class I-presented peptides in both melanoma, a “hot tumor” with high TMB, and EGFR mutant lung adenocarcinoma, a “cold tumor” with low TMB. We identified several classes of neopeptides, including mutated neoantigens and more than 1000 post-translationally modified peptides representing 58 different PTMs. We constructed a cancer germline (CG) antigen database with 285 antigens and identified 42 Class I-presented CG antigens. Finally, we developed a non-canonical peptide discovery pipeline to identify 44 lncRNA-derived peptides and validated Class I binding for select neopeptides. We provide direct evidence of HLA Class I presentation of a large number of neopeptides for potential vaccine or adoptive cell therapy in melanoma and mutant EGFR lung cancer.

## Introduction

Cancer immunotherapy has become one of the first line therapies for diverse cancers. The treatment outcome and patient survival rate are positively correlated with their tumor mutational burden (TMB) ^1, 2^. EGFR mutant lung adenocarcinoma occurs predominantly in never or oligo-smokers and exhibits a relatively low TMB ^3^. Immunotherapy has been less successful in EGFR mutant lung cancer, in part, because of its low TMB ^4, 5^. In contrast, melanoma, a cancer with high TMB due to UV exposure, responds well to current immune checkpoint blockade immunotherapy ^6, 7^. Consequently, the use of immunotherapy to treat low TMB cancers has been an unmet need. While classic immune checkpoint inhibition generally releases the natural endogenous immune response against cancer, more recently some success has been seen with adoptive T cell therapy (ACT) that creates a repertoire against “non-self” neoantigens or tumor associated antigens (cancer germline antigens) ^8–10^. Thus, the identification of cancer-specific or - associated antigen-derived peptides is important for the development of immunotherapeutic strategies for the treatment of “cold” tumors.

Recently, mass spectrometry (MS)-based proteomics has become a powerful approach for large-scale profiling of the human leukocyte antigen (HLA) Class I-associated peptidome ^11–13^. Unlike traditional HLA-epitope prediction algorithms, MS sequencing provides direct experimental evidence of the presented peptides, and it allows for the relative quantification of cell surface peptide presentation ^14^. This high throughput method can be used to profile thousands of *in vivo* HLA-associated immunopeptides ^15^. When combined with next-generation sequencing (NGS), to reveal somatic mutations, this approach is capable of detecting mutant neopeptides^16^ and non-canonical peptides derived from noncoding regions^17^. However, despite the fact that MS-based cancer antigen discovery has been widely employed for directly assessing antigen presentation, many previous studies only focused on “hot” tumors^18^ (e.g., melanoma, and high TMB lung cancers). Here, our goal was to develop a comprehensive proteogenomic platform to identify potentially targetable Class I-presented peptides in both melanoma, a “hot tumor” and lung adenocarcinoma, a “cold tumor”. We hypothesized that both low TMB-associated EGFR mutant lung tumors and high TMB-associated melanoma present a repertoire of tumor-associated or -specific peptide antigens on HLA Class I.

To develop this method, we enriched the cell surface presented HLA Class I bound peptides in two primary melanoma cell lines, two EGFR mutant lung adenocarcinoma cell lines and one primary tumor from an EGFR mutant patient who had undergone EGFR tyrosine kinase inhibitor (TKI) therapy. The high-resolution MS data of the HLA peptidome was searched against a comprehensive database generated from combining the normal human proteome as well as sample/patient-specific predicted proteome derived from NGS-identified mutations. Additionally, we leveraged deep learning-based *de novo* searching to deconvolute the remaining unmatched peptides. Our datasets contain five major clusters of potential cancer antigen sources; 1) common driver oncogenes, 2) PTMs, 3) cancer germline antigens, 4) mutated neoantigens, and 5) long non-coding RNAs (lncRNAs). We also selectively validated our findings utilizing targeted proteomics and an HLA stability binding assay. This study, for the first time, profiles potentially actionable HLA Class I-presented T cell epitopes in the immunologically “cold” EGFR mutant lung cancer that can be utilized for precision immunotherapy.

## Results

### Experimental design and identification of HLA Class I-associated immunopeptides

To test whether a large number of tumor antigen-derived peptides can be characterized in low TMB tumors, we selected two EGFR-mutant lung adenocarcinoma cell lines, PC9 and H1975, harboring EGFR^Del E746-A750^ and EGFR^L858R/T790M^, respectively. We also utilized a primary lung tumor harboring EGFR^L858R^, procured at autopsy from a patient (NCI-RA007) treated with the third generation EGFR tyrosine kinase inhibitor, osimertinib. In addition, two melanoma cell lines, NCI-3784Mel and NCI-3795Mel, generated from tumors of melanoma patients treated at the NIH Clinical Center, that represented tumors with high TMB. We employed high resolution MS-based peptide sequencing, NGS of genomic DNA/RNA and computational algorithms to identify 35,233 HLA Class I-associated peptides containing 8 to 14 amino acid residues. These include 14,876 database-searched peptides and 20,357 *de novo* searched peptides **(Supplementary Table 1)**. We identified somatic mutations by whole exome sequencing (WES) of tumor or tumor-derived cell lines and corresponding germline DNA from patients NCI-RA007, NCI-3784Mel and NCI-3795Mel. Expressed somatic mutations were also identified by RNA sequencing (RNA-seq) of the cell lines and tumor. PC9 and H1975 lung adenocarcinoma cell lines did not have corresponding germline DNA, hence all mutations identified by WES and RNA-seq were included for our data analyses. The mutations identified included single nucleotide variations (SNVs), small insertions-deletions (INDELs) and fusions. The HLA Class I antibody purified cell surface HLA binding peptides were sequenced by high-resolution tandem MS. The mass spectrometry data was searched against cell line or tumor-specific databases created by adding all corresponding mutant peptides to the normal human database (see Methods). The MS data was also searched using the *de novo* search algorithm in PEAKS studio (**Figure 1a** **and Supplementary Figure 1a**). As expected, EGFR-mutant lung adenocarcinoma patient NCI-RA007 had much fewer somatic mutations (289) compared to the two melanoma patients-derived cell lines, NCI-3784Mel and NCI-3795Mel (2678 and 2031, respectively) (**Supplementary Figure 1b** and **Supplementary Table 2**). Since HLA proteins are highly polymorphic and their binding peptides are HLA allele-restrictive ^19, 20^, HLA serotyping is important to further characterize the HLA Class I -associated peptides identified. We HLA typed the tumors and cell lines using seq2HLA analysis of the WES data ^21^. It is interesting to note that NCI-3784, H1975 and NCI-RA007 had loss of heterozygosity (LOH) of *HLA-B* and/or *-C* alleles (**Table 1**). HLA LOH has been suggested as a mechanism of immune escape ^22^. The total HLA Class I protein expression detected by immunoblotting (**Supplementary Figure 1c**) was consistent with cell surface HLA Class I presentation analyzed by flow cytometry (**Supplementary Figure 1d**). We performed 3∼6 biological replicates of HLA Class I-pull down experiments and MS analyses of associated peptides from each cell line/tumor. Pairwise correlation coefficients of peptide intensities from three representative biological replicates from PC9 cells show high correlation between replicates (**Figure 1b**). We identified 2385∼4401 database-matched peptides in each sample (**Figure 1c**). A majority of enriched immunopeptides were 9mer, consistent with the length of HLA class I bound immunopeptidomes reported previously^14, 23^ (**Figure 1d**).

**Figure 1.**
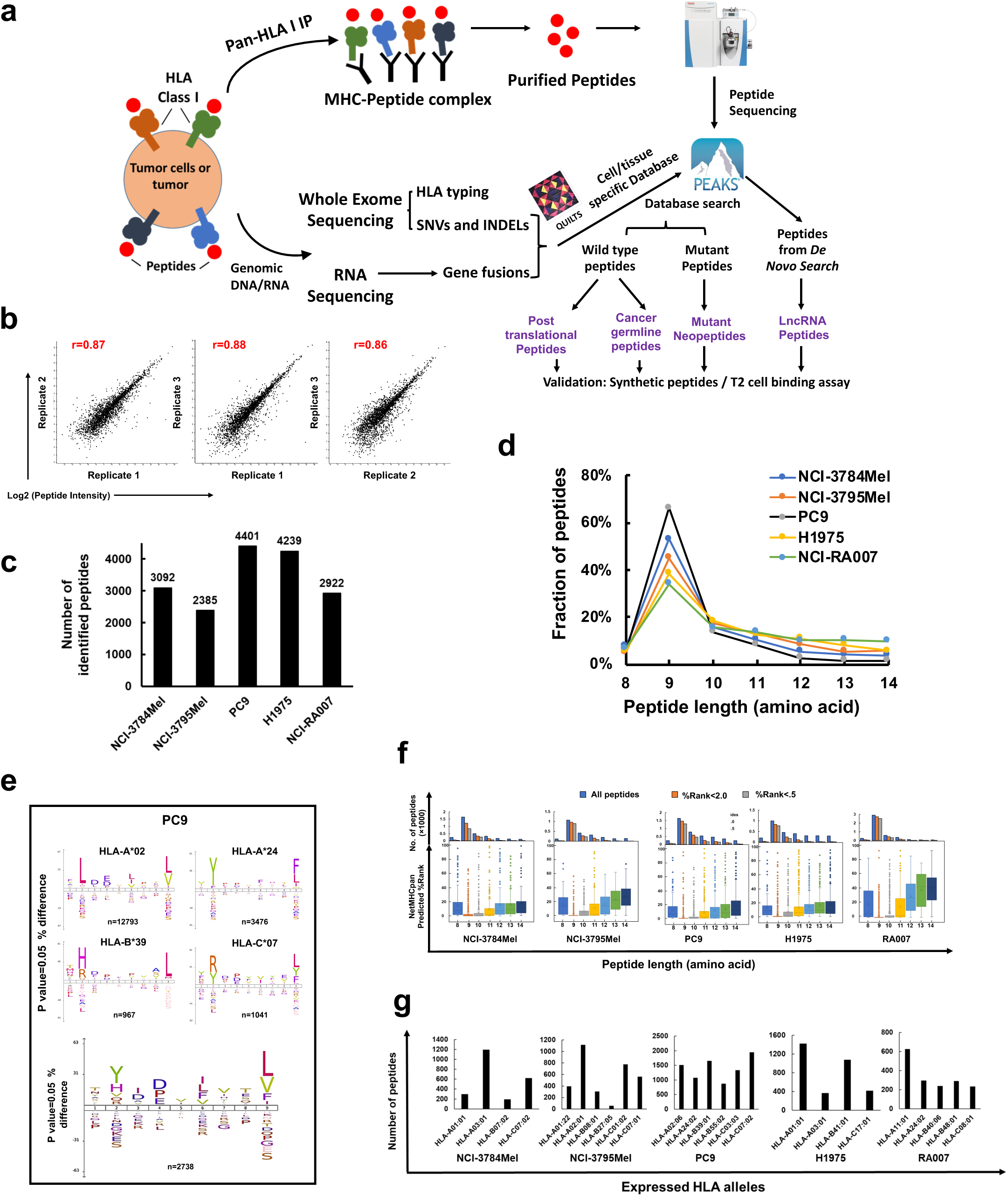
Overall proteogenomic strategy and identification of HLA Class I-associated immunopeptides from EGFR mutant lung cancer and melanoma patient-derived cell lines by mass spectrometry (MS). **a)** Overview of proteogenomic analysis pipeline for HLA immunopeptidome using MS-based proteomics and next generation sequencing (NGS). HLA Class I peptides were purified from two melanoma primary cell lines, two lung adenocarcinoma cell lines and a lung adenocarcinoma tumor and analyzed by high-resolution tandem MS. **b)** PC9 HLA Class I-associated peptide log2 transformed intensity identified by MS, and the Pearson’s correlation coefficients show high correlation among three biological replicates. **c)** Total number of peptides identified in the Class I immunopeptidomes from patient-derived melanoma and EGFR mutant lung cancer cells lines and tumor. **d)** The peptide length distribution within the Class I immunopeptidome from all samples shows that 9mer peptides are the most abundant. **e)** IceLogo motif analysis of four monoallelic-restricted 9mer peptidomes (HLA-A*02, A*24, B*39 and C*07) retrieved from experimentally validated IEDB database (upper panel) and MS-identified 9mer peptidome (lower panel) in PC9 cells. Percent difference stands for the % difference in frequency of the amino acids at a location **f)** NetHLApan 4.0 prediction algorithm-based scoring of each MS-identified peptide and distribution of binding scores among 8-14mer peptides. Upper panel shows the distribution of total identified peptides, binders (%Rank<2.0) and strong binders (%Rank<.5). Lower panel shows box plots of the lowest predicted %Rank for the corresponding HLA Class I allele for each peptide length. **g)** Number of predicted binders (%Rank<2.0) assigned to different HLA alleles for each sample based on Seq2HLA typing results.

**Table 1:**
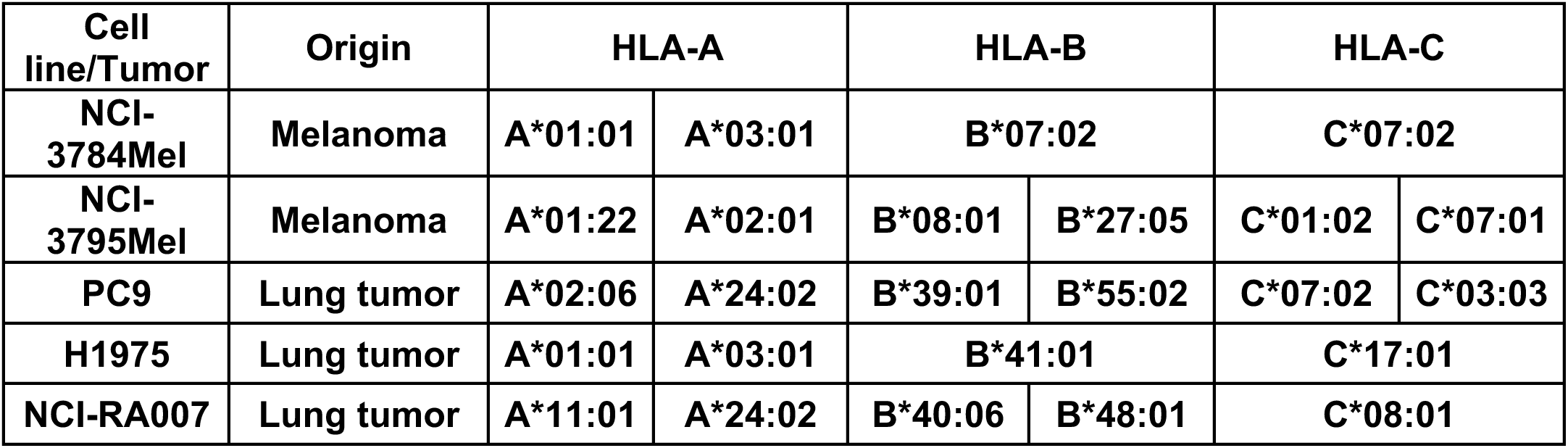
HLA tying of cancer cell lines and lung adenocarcinoma tumor.

| **Cell line/Tumor** | **Origin** | **HLA-A** |  | **HLA-B** |  | **HLA-C** |  |
| --- | --- | --- | --- | --- | --- | --- | --- |
| **NCI-3784Mel** | **Melanoma** | **A\*01:01** | **A\*03:01** | **B\*07:02** |  | **C\*07:02** |  |
| **NCI-3795Mel** | **Melanoma** | **A\*01:22** | **A\*02:01** | **B\*08:01** | **B\*27:05** | **C\*01:02** | **C\*07:01** |
| **PC9** | **Lung tumor** | **A\*02:06** | **A\*24:02** | **B\*39:01** | **B\*55:02** | **C\*07:02** | **C\*03:03** |
| **H1975** | **Lung tumor** | **A\*01:01** | **A\*03:01** | **B\*41:01** |  | **C\*17:01** |  |
| **NCI-RA007** | **Lung tumor** | **A\*11:01** | **A\*24:02** | **B\*40:06** | **B\*48:01** | **C\*08:01** |  |

Computational HLA binding epitope prediction algorithms have been widely recognized as a powerful tool to estimate HLA-ligand binding affinity and tumor neoantigen prediction ^24, 25^. To predict the HLA Class I restriction of the HLA Class I-associated peptides identified in our dataset, we extracted the customized mono-allelic experimentally validated binding peptidome from the Immune Epitope Database and Analysis Resource (IEDB) ^26^. Since 9mer is the most common length for peptides in the HLA Class I immunopeptidome, we used iceLogo to visualize the 9mer peptide-binding motifs of our enriched HLA Class I peptidome from tumor cell lines/tissue and their corresponding mono-allelic datasheets from IEDB^27^. For example, HLA-peptide binding motifs of four major HLA typed alleles in PC9 cells are the main components of endogenous enriched PC9 immunopeptide by overlaying the four-individual mono-allelic binding motifs (**Figure 1e**). Our data confirmed the Class I anchor positions (+2 and last residue) are highly conserved. We further analyzed our dataset using NetMHCpan-4.0^28^ and determined weak and strong binding using %Rank < 2.0 and <0.5, respectively. A majority of the enriched peptides were predicted to be binders; nonetheless, 9mer and 10mer peptides had lower scores and hence predicted stronger binding (**Figure 1f**). The predicted HLA-binders were assigned to the expressed HLA alleles in each sample. Notably, although A*01:01 and A*03:01 were both present in NCI-3784mel and H1975, they barely share any binding peptides, suggesting the possibility of different HLA ligand processing and presentation machinery in melanoma and lung cancer **(Supplementary Figure 2a-b)**. We observed a similar phenomenon in HLA-A*24:02 (typed in PC9 and NCI-RA007) and Cw*07 (typed in PC9, NCI-3784Mel and NCI-3795Mel) (**Supplementary Figure 2c-d**).

### Functional annotation of Class I-associated peptide parent proteins

We classified all presented immunopeptide source proteins according to their subcellular localization (**Figure 2a**) and molecular function (**Figure 2b**), and discovered that the endogenous immunopeptidome source proteins and cellular proteomes had similar trends of distribution ^29^. This suggests all intracellular proteins may generate HLA Class I-presented peptides. Utilizing the MS-detected intensity of the immunopeptides as a surrogate for parent protein expression levels, and a cellular functional bioinformatic analysis tool (www.proteomaps.net ^30^), we found significantly enriched proteins are involved in metabolic processes, such as oxidation, phosphorylation and amino acid metabolism, cell cycle, ubiquitin labeling, proteasome regulation, MAPK signaling, transcription, spliceosome and RNA transport (**Figure 2c**). Ingenuity Pathway Analysis (IPA) ^31^ of identified parent proteins indicated that EIF2, EIF4,mTOR, sirtuin, integrin, glucocorticoid receptor, PI3K and actin signaling pathways, ubiquitination and phagosome maturation pathways were enriched. The enrichment score indicates the difference of source protein a) between lung adenocarcinoma and melanoma and b) between cell lines and tumor tissue. (**Figure 2d**). IPA upstream regulator analysis showed tumor suppressor TP53 and proto-oncogenes *MYC*, *KRAS*, *ESR1*, *ERBB2*, *EGFR, RICTOR* and *mTOR* to be among the top upstream potential regulators of the parent proteins identified (**Figure 2e**). Network analysis confirmed that tumor suppressors (e.g., TP53, BRCA1) and oncogenes (e.g., EGFR, KRAS, MYC) were components of the network of parent proteins represented by the identified Class I-associated peptides (**Figure 2f**). To further supplement the bioinformatics analyses, we identified HLA Class I -presented peptides from common protooncogenes, such as *KRAS*, *EGFR*, *MYC*, *JUN*, and tumor suppressors, such as *TP53*, *RB1* and *BRCA2*. We identified 17 common cancer driver gene-derived wild type peptides presented by HLA Class I in lung adenocarcinoma cell lines/tumor and 2 peptides in primary tumor-derived melanoma cells; 6 of which are novel peptides not previously reported (**Figure 2g**). In addition, we used NetMHCpan to predict the HLA Class I restriction of the identified cancer driver gene-derived peptides. We found that, though some peptides have been predicted to be binders for multiple respective HLA alleles (e.g., KQFEGTVEI derived from BRCA2), their MS intensity is not significantly higher than that of the peptides predicted to be monoallelic binders (e.g., KLISEEDLLRK derived from MYC) (**Supplementary Table 3**). Taken together, our analyses identified 19 high-confidence oncogene/tumor suppressor-derived epitopes with potential relevance to cancer immunotherapy.

**Figure 2.**
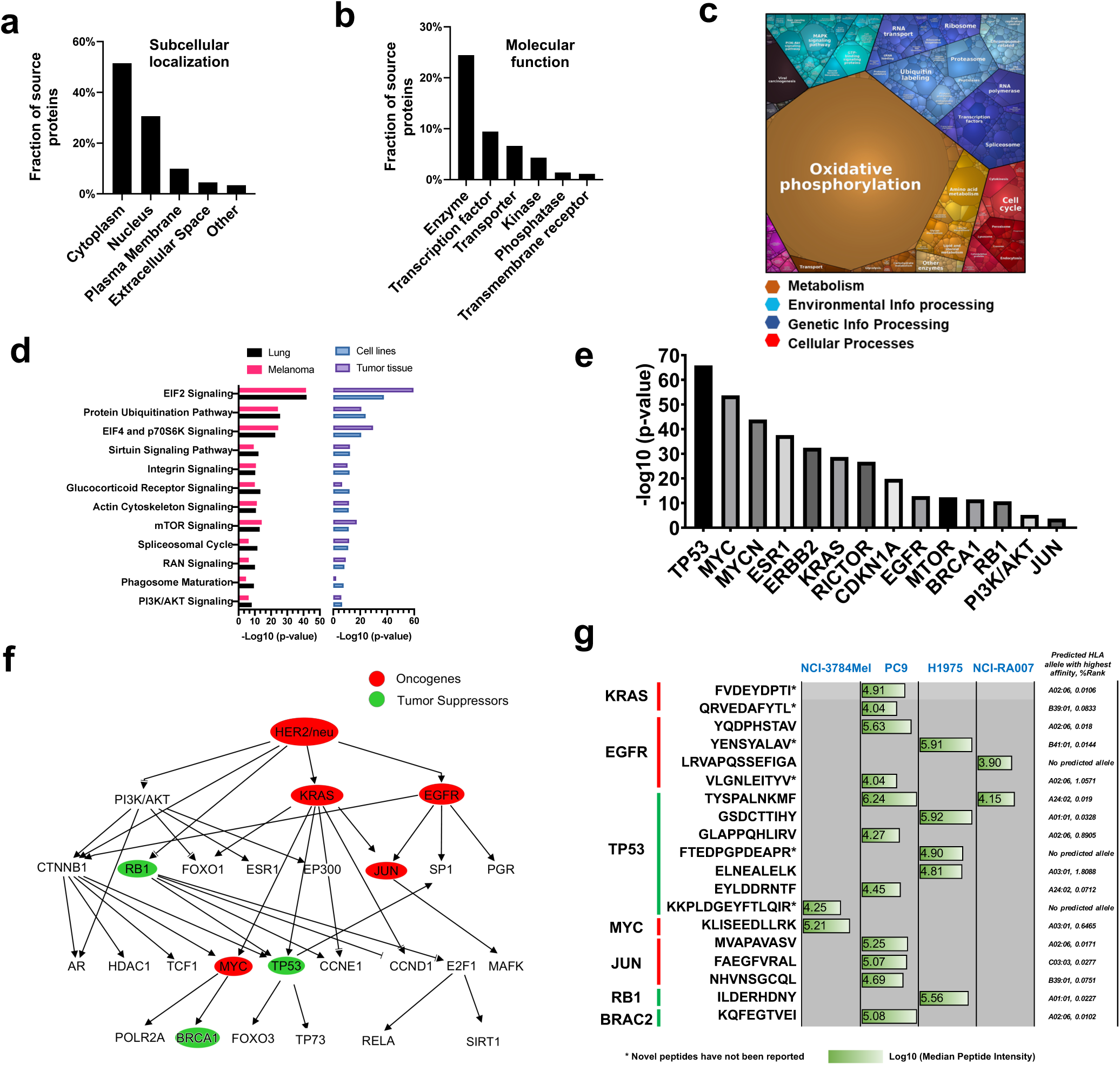
Functional annotation of HLA Class I -associated peptide parent proteins. **a)** Subcellular localization and **b)** molecular function classification of total identified HLA Class I-associated immunopeptides using Ingenuity Pathway Analysis. **c)** Proteomap analysis (www.proteomaps.net) of total identified HLA Class I-associated peptide source proteins, considering the annotated protein molecular function and abundance. **d)** Canonical pathways enriched among the identified HLA Class I-associated immunopeptide parent proteins which are grouped into either lung adenocarcinoma (i.e., PC9, H1975, NCI-RA007) and melanoma (i.e., NCI-3784mel and NCI-3795Mel) or cell lines (i.e., PC9, H1975, NCI-3784mel and NCI-3795Mel) and tumor tissue (i.e, NCI-RA007). **e)** Oncogenes and tumor suppressors, predicted by upstream regulator analysis, to be key regulators of the HLA immunopeptide source proteins. **f)** Network analysis of the predicted key upstream regulators of the of Class I peptide source proteins. **g)** Key oncogene and tumor suppressor Class I peptides identified in this study. Peptides have not reported to date as Class I presented are indicated with *.

### *In vivo* post-translationally modified (PTM) peptides are presented by HLA Class I and are potential neoantigens

To identify the *in vivo* PTM peptides, we used PEAKS studio to search the MS raw files against the UniProtKB human database, using the pan-PTM option (over 650 variable modifications) at stringent 1% FDR. Peptide modification artifacts induced by sample preparation (e.g., urea, reducing and alkylating reagents) and electrospray ionization (ESI) are the major concerns for *in vivo* PTM identification ^32^. Our HLA Class I-associated peptidome enrichment protocol did not involve the use of urea buffer, protein reduction and alkylation or enzyme digestion. ESI artifacts could be excluded by examining the retention time of modified peptides and their unmodified counterparts; artifact modifications were only added during the ionization so those modified peptides must be co-eluted with their unmodified counterparts, yet we did not observe co-eluted peptides. We identified 1351 modified and 11841 unmodified 8-14mer peptides. We did not detect the corresponding unmodified form for 804 of the modified peptides, this group of peptides are defined as “modified only”. On the other hand, for 411 of the unmodified peptides we identified 543 modified counterparts (this is a group of peptides defined as “modified and unmodified”, suggesting the existence of multiple PTMs for some peptides (**Figure 3a** **and Supplementary Table 4**). Approximately 10% of the total peptides identified had at least one PTM, and we identified 58 different PTMs making this the largest HLA PTM immunopeptidome identified to date. The heatmap shows the median intensity of each PTM in each sample whereby methionine oxidation (639 peptides), deamination (176 peptides), acetylation (144 peptides), and methylation (78 peptides) were the most abundant modifications seen (**Figure 3b**), in agreement with previous reports^14, 15^. Interestingly, the length distribution of the PTM peptides was quite different from the classic the HLA Class I bound peptidome profile; where more peptides were longer than 9 amino acids. It is possible that the peptide conformation has been altered by the PTM, especially when the modifications happen on the HLA molecule anchor positions. (**Figure 3c**). Interestingly, among the 9-mer peptides identified, the N-terminal amino acid was most commonly modified; the first amino acid was modified in 102/293 9-mer PTM peptides (**Figure 3d**). PTM HLA peptides were previously shown to be more abundant than their unmodified counterparts^15^. However, in our dataset pan-PTM peptides were significantly less abundant (as measured by intensity) than unmodified peptides (p-value=5.4E-16) (**Figure 3e**). Similarly, MS intensity was lower for deaminated and methylated peptides than unmodified peptides (**Figure 3f-g**). However, the median intensity of glutamate to pyroglutamate (pyroGlu) modified peptides was similar to that of their unmodified counterparts (**Figure 3h**). The pyroGlu modification occurred on gultamate at N terminal position one, which does not significantly affect the peptide conformation and binding affinity^33^.

**Figure 3.**
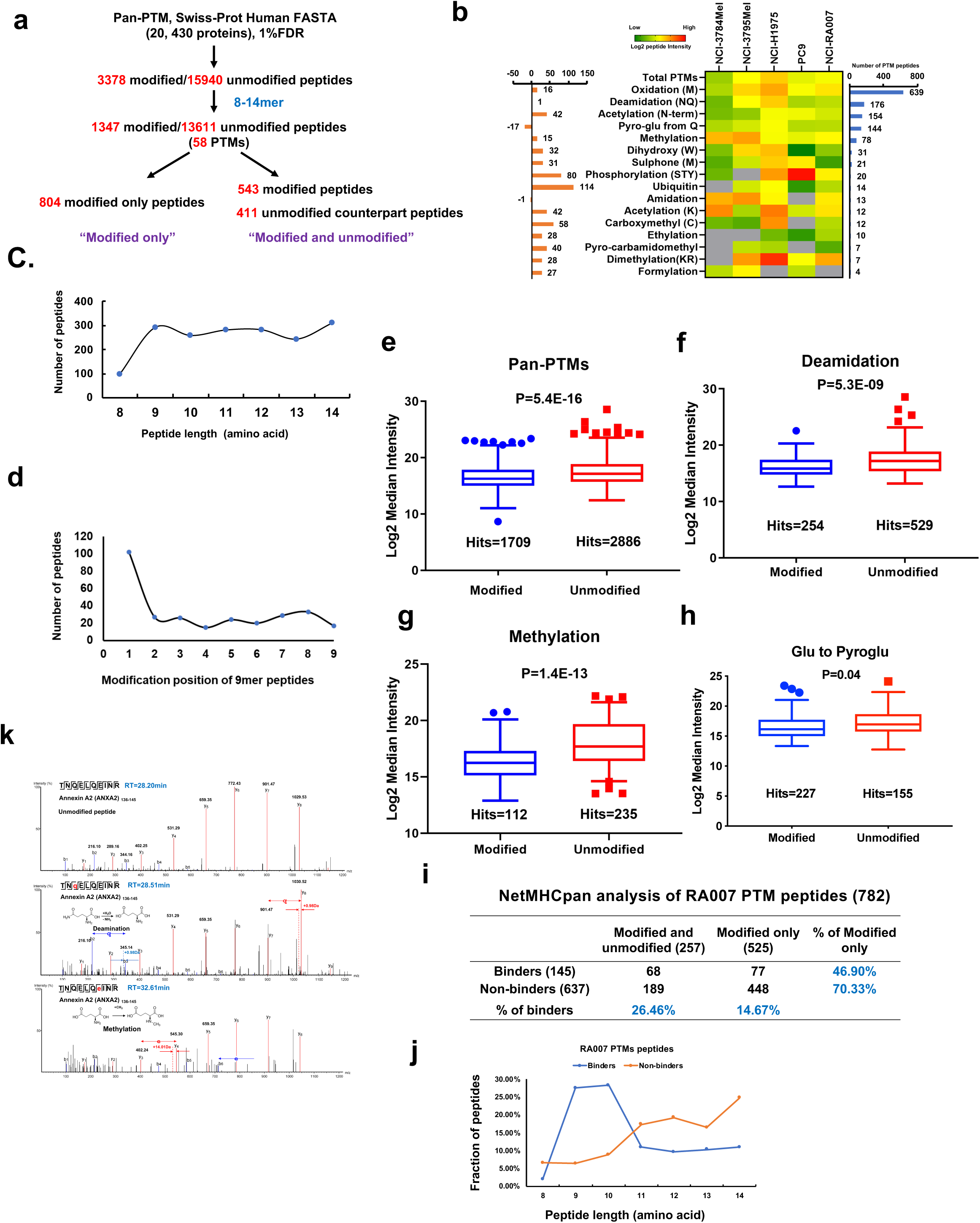
Characterization of pan-posttranslationally modified Class I -associated immunopeptides. **a)** Pan-PTM-derived immunopeptidome profiling pipeline. **b)** Heatmap shows the median log2 intensity of all identified PTM peptides; left bar graph displays the molecular weight shift of each PTM; right bar graph shows the total number of peptides identified in each PTM group. **c)** Length distribution of PTM peptides. **d)** Post-translationally modified amino acid position distribution of 9mer modified peptides. **e-h)** Box plots showing the median log2 intensities of modified peptides (blue) and their unmodified wt counterparts (red) for: **e)** pan-PTM **f)** Deamidation **g)** Methylation **h)** Glutamate to Pyroglutamate conversion (Glu to Pyroglu). **i)** NetMHCpan analysis of 782 PTM peptides identified in sample NCI-RA007 where peptides with %Rank<2.0 are considered binders. Table shows the percentage of predicted “binders” in the “unmodified and modified” and “modified only” peptides; it also shows the percentage of “modified only” peptides in the “binders” and “non-binders” peptides. **j)** Peptide length distribution of predicted binder and non-binder PTM peptides identified in RA007. **k)** MS/MS spectra of one representative peptide with multiple PTMs. ANX2-dervied peptide TNQELQEINR, identified as unmodified (top panel), deaminated (middle panel) and methylated (lower panel).

To validate the data quality and verify that the modifications were generated *in vivo* and not ionization artifacts, we selectively examined the tandem mass spectra of the modified peptides. A deaminated and a methylated form of peptide TNQELQEINR, derived from Annexin A2 (ANXA2), had a retention time (RT) of 28.51min and 32.61min, respectively, whereas the unmodified peptide had a RT of 28.20min (**Figure 3k**). We manually verified the RT of methionine oxidized peptides and their unaffected counterparts and determined that they did not co-elute during LC. Herein, we can conclude that the PTM peptides identified were generated *in vivo*.

Next, we determined whether the PTMs may alter the binding affinity to HLA Class I. There are no HLA binding prediction algorithms commercially available that accounts for PTM peptides. Using the unmodified forms of the peptides identified for NetMHCpan4.0 analysis of RA007, 637 of 782 PTM peptides were considered to be non-binders (%Rank>2.0), suggesting that specific modifications of these peptides may have been critical for HLA binding. Notably, the percentage of predicted binders for peptides with both modified and unmodified counterparts was nearly twice (26.46%) that of solely modified (14.67%). Also, the percentage of predicted binders for modified only peptides (46.90%) was much lower than that of non-binders (70.33%) (**Figure 3i**). We plotted peptide length distribution of the binders and non-binders and found that they had a similar pattern although more non-binders were longer peptides (>10mer) (**Figure 3j**). This further suggests that PTM peptides may have non-conventional length distribution for HLA Class I binding. Collectively, our findings suggest that PTMs may be important for a subset of HLA Class I-binding peptides and antigen presentation. More importantly, these peptides are more likely to be predicted as “non-binders” to HLA Class I using existing bioinformatic algorithms.

### Novel cancer germline (CG) antigen-derived peptides expressed in melanoma and lung adenocarcinoma

Cancer germline (CG) or tumor testis antigens hold great potential for generating tumor specific antigens for T-cell-based therapy^34^. CG antigens are exclusively expressed or overexpressed in tumor and germ cells. These peptide antigens are rarely presented to immune cells due to the relatively low HLA expression in testis and germline tissues^35, 36^. In order to search for these CG antigens in our mass spectrometry data for Class I molecule-associated peptides, we established a customized CG antigen library by compiling 285 CG antigens using the CTDatabase (http://www.cta.lncc.br/), Human Protein Atlas (https://www.proteinatlas.org/) and Cancer Antigenic Peptide Database (https://caped.icp.ucl.ac.be/), and literature searches (**Figure 4a** and **Supplementary Table 5**).The 20,430 Class I-associated peptides identified by MS and annotated by a UniProtKB database search at 1% FDR were next queried against this CG antigen library. We identified a total of 42 CG antigen peptides derived from 14 CG antigen proteins. We then used synthetic peptides and targeted proteomics to further validate nine of these peptides from the melanoma patient-derived cell lines (and **Supplementary Table 6**). Of the 42 CG antigen-derived peptides, 29 are from melanoma and 12 from lung adenocarcinoma, and of these 10 were novel and, to our knowledge have not been reported previously (**Figure 4b**). To validate the peptides, we visualized and inspected MS2 spectra of each peptide manually. Furthermore, we validated the nine peptides and one novel peptide identified in melanoma cells and NCI-RA007 tumor, respectively, by matching the mass spectra of synthetic peptides (purity > 99%) (**Figure 4c** **and Supplementary Figure 3b-c**). We identified 15 melanocyte protein PMEL (GP100)-derived epitopes in the 3784Mel cell line derived from the tumor of a patient whose tumor-infiltrating CD8^+^ lymphocytes (TILs) have been previously shown to recognize the GP100 antigen ^37^. Since CG antigens have been extensively studied in melanoma, we only found one novel peptide, VTPVEVHIGT derived from sperm-associated antigen 17 (SPAG17). In contrast, and of particular interest, we identified seven novel peptides in the H1975 lung adenocarcinoma cell line. For example, these include peptides mapping to lactate dehydrogenase C (LDHC) and testis-expressed protein 15 (TEX15), Next, we verified whether the genes for these proteins were expressed at the transcript and protein levels in H1975 cells where the total RNA and total proteome were profiled separately. We mapped, quantified and normalized H1975 RNA-seq FASTQ files to reads per kilo base per million mapped reads (RPKM) and used MS-based proteomics to extensively fractionate profile and quantitate proteins extracted from H1975 cells. Interestingly, expression of TEX15 and LDHC RNA was lower and protein was not detected, underscoring the possibility that Class I presentation can occur for genes expressed at low levels (**Figure 4d**). We also ranked CG antigen gene and protein expression in H1975 cells and found representation of Class I-associated peptides for genes with protein levels undetectable by MS but detectable at the transcript level (**Figure 4e**). Therefore, our data supports the phenomenon that immunopeptidome lacked association with gene/protein expression^14^.

**Figure 4.**
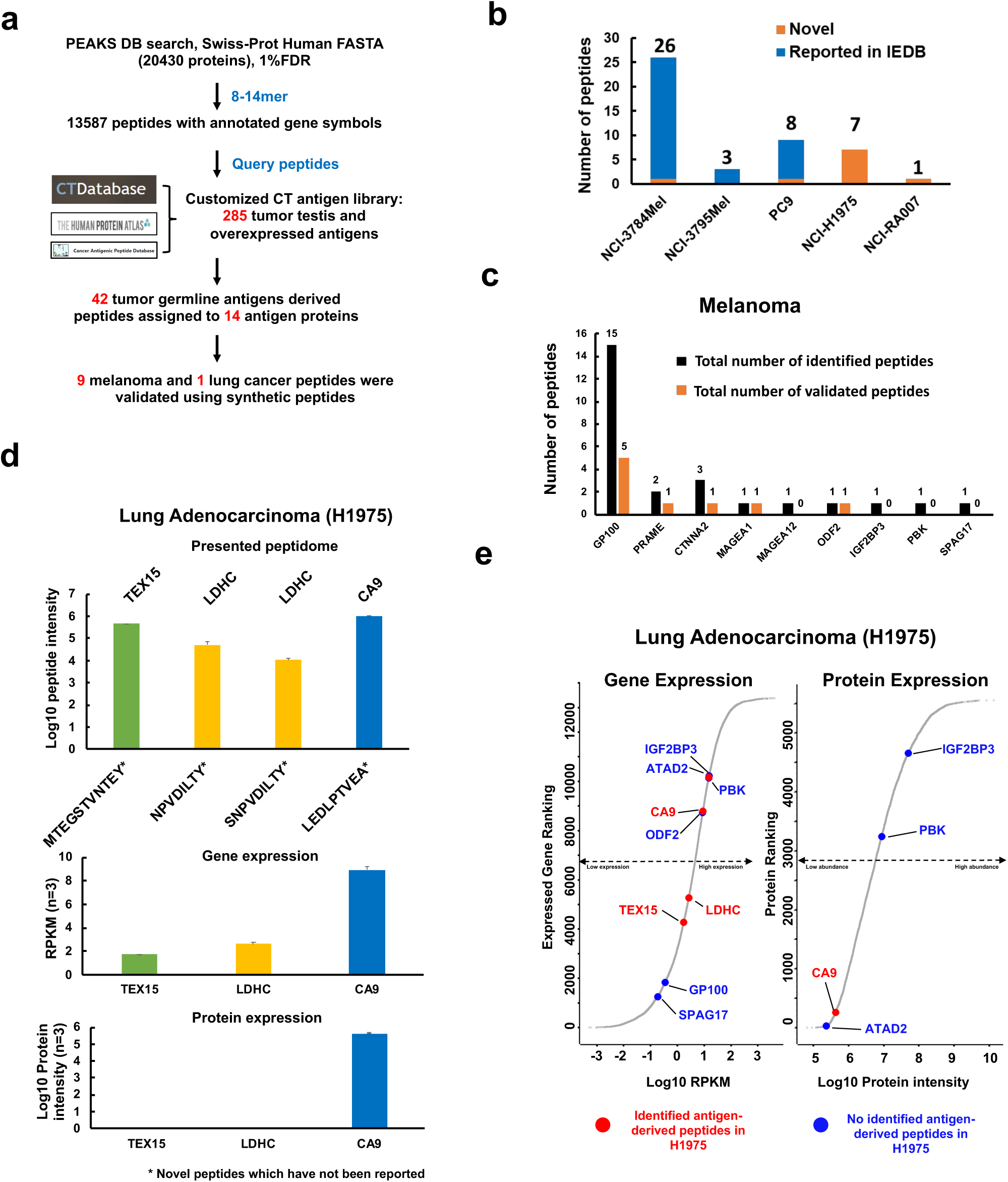
Identification of cancer germline (CG) antigen-derived peptides. **a)** Overall strategy of CG antigen identification using customized library integrating reported immunogenic and manually added CG antigens based on their expression level in different organs; a total of 285 CG antigens were included in this custom database. **b)** The number of novel and reported CG antigen-derived peptides. **c)** 28 peptides identified in NCI-3784Mel assigned to 9 CG antigens. **d)** Log10 median peptide intensity of 4 immunopeptides identified in H1975 (top panel); gene expression (middle panel) and protein expression (lower panel) of 3 source antigen proteins. **e)** Ranking of gene and protein expression of CG-antigens identified in H1975. Red dots indicate the antigen protein-derived peptides have been identified in H1975, and the blue dots indicate the antigen protein-derived peptides haven’t been identified in H1975.

### Lung adenocarcinoma neopeptides are potential neoantigens for cancer immunotherapy

Mutant neopeptides derived from somatic mutations in tumors, if presented by MHC Class I, have the potential to engage cytotoxic T cells to promote tumor immunity. Hence the identification of mutant neopeptides presented by the cognate Class I proteins is of paramount importance. To identify mutant neopeptides, we first constructed cell line and patient-specific databases by adding all somatic variants identified by NGS (RNAseq and WES) to normal human database used for searching MS data (**Figure 5a**) when germline DNA was available for sequencing (NCI-3784Mel, NCI-3795Mel and NCI-RA007). For PC9 and H1975 cells which do not have available germline DNA, all variants identified by exome sequencing were used. We identified 12 peptides harboring SNVs in the two lung adenocarcinoma cell lines; but no INDELs and fusions. Further analyses of these variant neopeptides using NetMHCpan predicted all to be binders for at least one HLA allele in their corresponding cell line. Pathogenic variant peptides were identified using COSMIC, dbSNP and ClinVar (NCBI). Population-based mutant allele frequency indicates that 4 neopeptides (minor allele frequency<0.05) are potential neoantigen-derived targets, and 8 neopeptides (minor allele frequency>0.05) could result from the single nucleotide polymorphisms (SNPs) (**Figure 5b**). We compared the MS intensities of the 12 mutant neopeptides and the non-mutated Class I -associated peptides and found no significant difference in median peptide intensities between the two groups for both the H1975 and PC9 cell lines (**Figure 5c-d**), suggesting significant Class I presentation of the mutant neopeptides. In addition, to validate the neopeptide sequences identified in our Class I pull-down experiments, we synthesized matching peptides and utilized LC tandem MS to compare both sets. In fact, 11 out of 12 synthetic peptides co-eluted with the endogenous neopeptides and their MS2 spectra were an exact match, two examples of which are shown in **Figure 5e-f**. Next, we confirmed the binding of the neopeptides to specific predicted HLA alleles. Peptide SIT<u>S</u>IISSV, derived from RIF1p.G836S binds strongly to HLA-A*02:06 with %Rank at 0.23. To assay cell surface HLA stabilization, we pulsed this peptide overnight to an antigen peptide transporter (TAP)-deficient T2 cell line that expresses only HLA-A*02 ^38^, and used HLA-A*02 binding peptide SLLMWITQC from NY-ESO-1 as a positive binding control in parallel. Both peptides significantly stabilized HLA to the cell surface in comparison to the DMSO control (**Figure 5g-h**). We identified 12 variant neopeptides in lung adenocarcinoma cell lines which we extensively validated by multiple methods. Taken together, these results suggest these variants are potential neoantigens for cancer immunotherapy.

**Figure 5.**
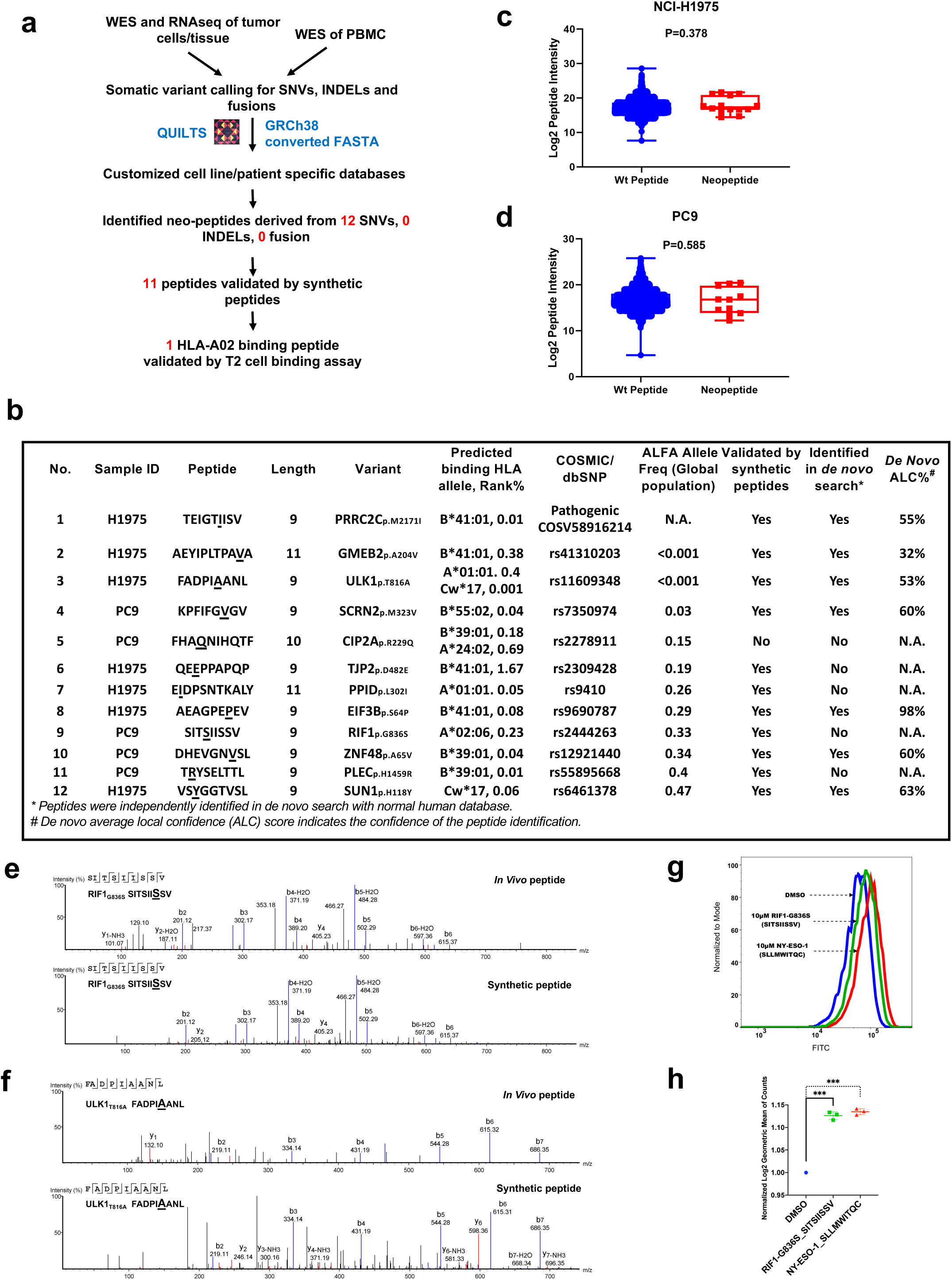
Identification of Class I -presented mutated neopeptides in lung adenocarcinoma. **a)** Workflow of integrated proteogenomic analysis using germline (PBMCs) and tumor cell line/tissue WES and RNAseq datasets to identify SNVs, INDELs and fusions and construction of tumor cell line/tumor tissue-specific databases to interrogate the MS data of Class I -associated peptides. **b)** List of 12 mutated neopeptides, with its variant, predicted HLA-allele restriction, dbSNP ID and synthetic peptide validation and *de novo* sequencing search status. **c-d)** Box plots show peptide intensity of wildtype and mutant nonpeptides in all biological replicates from **(c)** H1975 and **(d)** PC9. **e-f)** Matched MS2 spectra of endogenous and its synthetic counterpart for **(e)** RIF1-G836S-derived neopeptide SITSIISSV and **(f)** ULK1p.T816A-derived neopeptide FADPIAANL. **g)** T2 cell-based HLA stability assay shows that RIF1 p.G836S -derived peptide (SIT*<u>S</u>*IISSV) bound and stabilized HLA-A*02. **h)** Box plot shows statistically significant increase of HLA expression (log2 geometric mean of counts) in T2 cells incubated with RIF1 p.G836S -derived peptide (SIT*<u>S</u>*IISSV) and positive control NY-ESO1-derived peptide compared to those incubated with DMSO (p<0.005).

### *De novo* sequencing improves peptide identification compared to database search

*De novo* searching of MS spectra from large-scale mass spectrometry analysis has been employed by various algorithms, including PEAKS studio. The impressive prediction accuracy of this approach has been extensively reported ^39, 40^. We searched our entire Class I-associated immunopeptidome MS data using the PEAKS studio *de novo* search algorithm. First, to evaluate the data quality of the peptides identified by *de novo* sequencing, we manually inspected the MS2 spectra. We then employed the NetMHCpan prediction to evaluate the binding capacity of our *de novo* peptides. We determined that while an average of ∼55% of the database-searched (DB) peptides were predicted to be specific HLA allele binders for their corresponding cell line/tissue, an average of ∼33% of *de novo* peptides were predicted to be a strong binder of at least one HLA allele in the respective sample (**Figure 6a-b**). We observed a relatively robust correlation of peptide intensities between biological replicates (Pearson’s coefficient = 0.74) of peptides identified by *de novo* sequencing analysis of the MS data (**Figure 6c**). We employed a similar approach employed to evaluate the HLA binding affinity of 8-14mer peptides identified by DB search (**Figure 1f**) by predicting the binding affinity and assigning each peptide to its highest predicted HLA allele and showing the distribution of 8-14mer peptides according to their lowest %Rank (**Figure 6d**). We observed the same trend with respect to binding score distribution of 8-14mer peptides when comparing *de novo* to DB identified peptides. The 9-mer peptides have the lowest binding scores and hence highest binding affinity to the best predicted HLA binding allele (**Figure 6d**). We also assigned each peptide to its highest predicted HLA allele and estimated the total number of peptides assigned to each of the HLA alleles (**Figure 6e**). To further confirm the validity of the peptide identities from the *de novo* sequencing analysis, we compared the predicted DB and *de novo* 9-mer HLA-binding peptide motifs. Interestingly, the peptide motifs were very similar between the two groups, reinforcing the validity of the *de novo*-identified peptides generated from all 5 samples (**Figure 6f-j**). We further validated our *de novo* sequencing pipeline-identified mutated neopeptides from the lung adenocarcinoma cell lines by comparison to those identified using cell line-specific database search. Indeed, we found four of seven and two of five mutant neopeptides in H1975 and PC9 cells, respectively (**Figure 6k**). We next aligned the *de novo* and DB search spectra for one representative endogenous peptide (AEAGPEPEV) to that of the synthetic peptide. The b and y ions were perfectly aligned among the 3 spectra (**Figure 6l**). In summary, we present strong evidence that this is a robust and reliable *de novo* sequencing pipeline for MS identification of the immunopeptidome.

**Figure 6.**
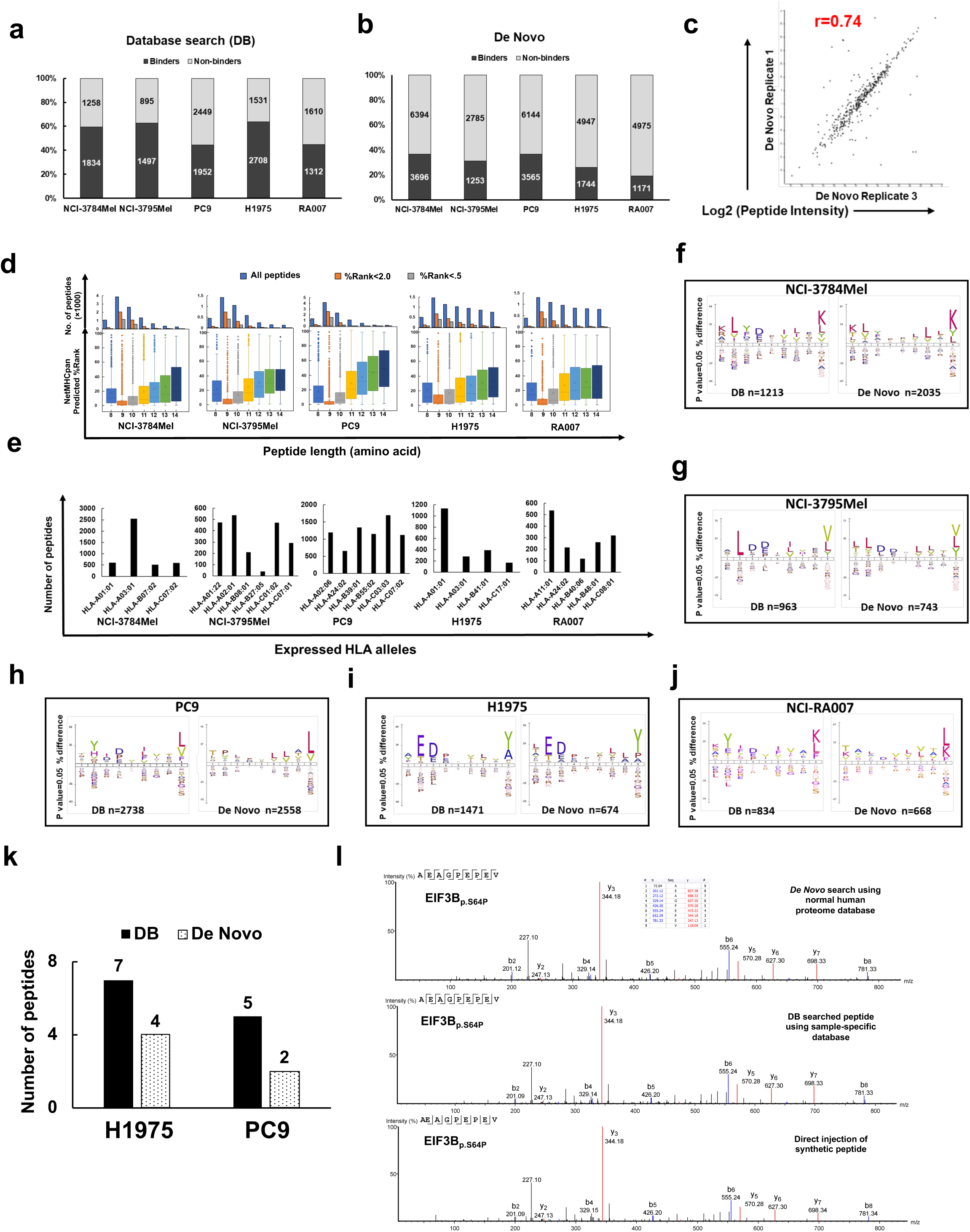
*De novo* search of MS data to identify Class I -associated immunopeptidome. **a-b)** Fraction of Class I -associated peptides predicted to be HLA binders (NetMHCpan %rank<2.0) or non-binders (%Rank>2.0) by **(a)** database search and **(b)** *de novo* sequencing. **c)** Correlation of peptide intensities from *de novo* sequencing algorithm-searched peptides from two PC9 biological replicates. **d)** Distribution of total number of *de novo* sequencing-searched 8-14mer peptides, HLA binders (%Rank<2.0) and strong binders (%Rank<0.5) (upper panel). Distribution of HLA binding affinity (%Rank) of *d*e *novo* sequencing-searched 8-14mer peptides (lower panel). **e)** Number of *d*e *novo* sequencing-searched binders assigned to different HLA alleles. **f-j)** Comparison of 9 mer peptide binding motifs (binders only, NetMHCPan %Rank<2.0) identified by database search (DB) search versus *d*e *novo* search in **(f)** NCI-3784Mel, **(g)** NCI-3795Mel, **(h)** PC9, **(i)** H1975 and **(j)** NCI-RA007. **k)** Of 12 mutated neopeptides identified by cell line/tumor specific DB search, six were also identified by *de novo* search. **L)** Matched MS2 spectra of one representative endogenous neopeptide, EIF3B p.S64P (AEAGPE*<u>P</u>*EV), identified by *de novo* search (upper panel), proteogenomic DB search (middle panel) and direct injection of its synthetic peptide (lower panel).

### Identification of MS-identified long non-coding RNA (lnc-RNA)-derived peptides using *de novo* sequencing

Non-coding regions in the genome are the most unexplored, yet rich source of neoantigens. Previous studies have profiled the non-coding immunopeptidome using the traditional proteogenomic approach of searching the MS raw files against sample-specific library generated from RNA-seq data ^41, 42^, which always resulted in extremely massive customized noncoding sequence libraries because 99% of the human genome is noncoding ^43^, a majority of which are unannotated. We developed a profiling pipeline of potential lncRNA-derived peptides by taking advantage of deep *de novo* analysis of MS data without using a pre-define database and then matching the MS-identified peptides with hypothetical peptides generated by 6-frame translation of all lncRNAs from an available lncRNA database, LNCipedia containing ∼50,000 lncRNAs from the high-confidence genome assembly ^44^. We queried the 8-14mer *de novo* peptides against all six potential reading frames of the translated LNCipedia-derived protein database. We also confirmed the transcript expression of the lncRNAs coding the identified peptides in RNA-seq gene expression data from the patient-derived cell lines. We used the Integrative Genomics Viewer (IGV) to visualize peptide coding regions of these lncRNAs. We finally validated the endogenous peptides derived from the lncRNAs with synthetic peptide-based targeted MS and T2 cell-based HLA stability assay for binding to specific HLA alleles (**Figure 7a** and **Supplementary Figure 4a**). A total of 195 distinct *de novo* sequencing-identified peptides matched to the six frame-translated lncRNAs in the LNCipedia database, of which, 71 were predicted to be binders (%Rank<2.0) for at least one HLA allele in their corresponding cell line/tumor. We further analyzed the RNA-seq data and found that the source RNAs of 53 peptides were transcribed. The FeatureCount intensities of these transcribed lnRNAs were displayed in a heatmap (**Supplementary Figure 4b**). We then confirmed 44 peptides had their specific coding regions transcribed and did not overlap with any protein-coding region using the BLAST-like Alignment Tool (BLAT) in the IGV (**Figure 7b** and **Supplementary Table 7**), two representative examples of data visualization are shown in **Supplementary Figure 4c-d.** Notably, we didn’t observe any peptide presented in more than two samples, implying that lncRNA-derived peptides may be tumor cell line and patient-specific. To assess the significance of our findings, we generated a mock lncRNA pool from randomly sampled ∼50,000 genomic sequences. We rejected the null hypothesis that our identified lncRNA-derived peptides randomly matched to six frame-translated LNCipedia-derived database, as we obtained a significant empirical p value (1.1e^-5^) upon comparing to the random matches against the mock lncRNA pool-derived 6 frame translated database.

**Figure 7.**
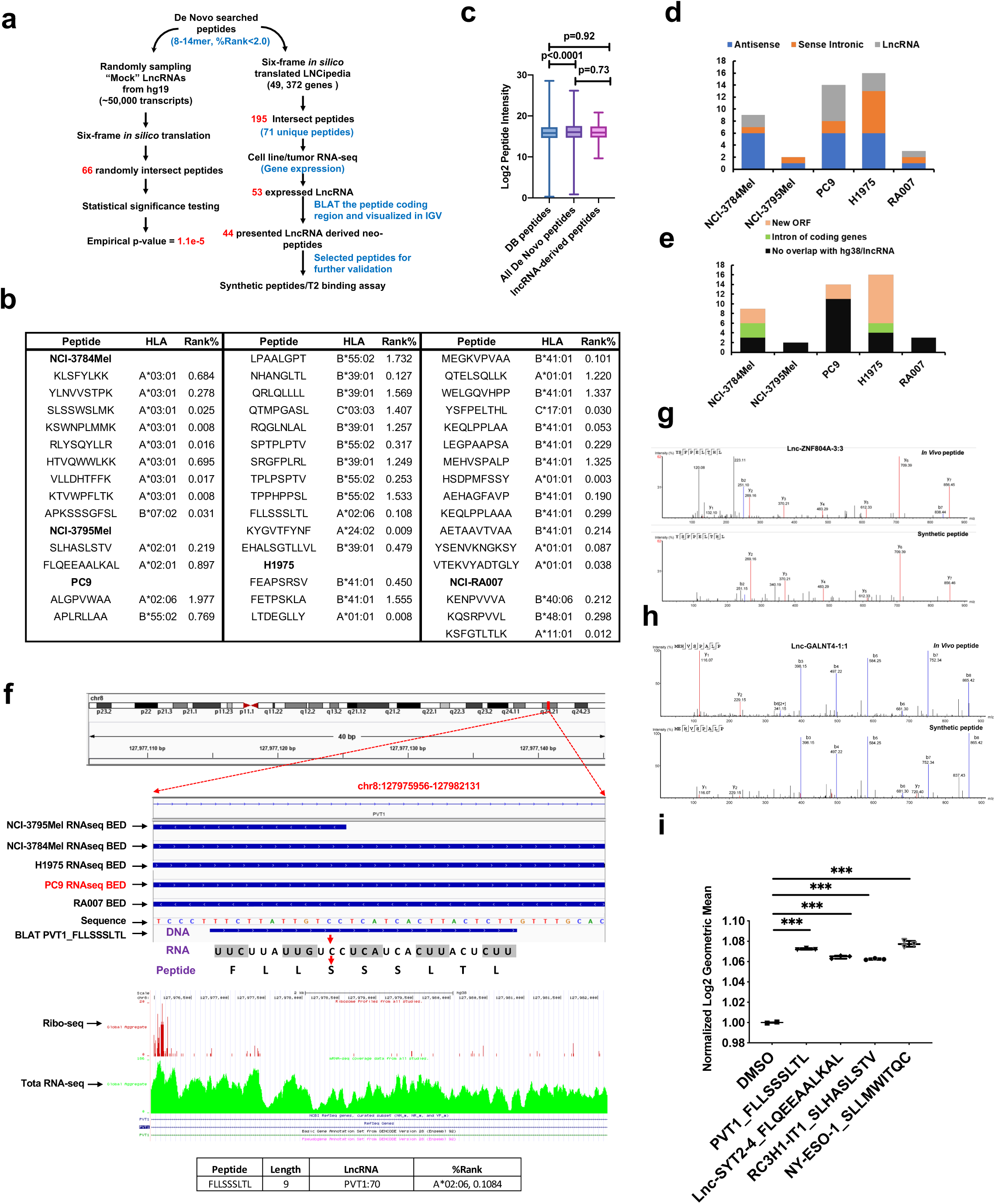
Identification of lncRNA-derived neopeptides by proteogenomic and *de novo* sequencing analyses. **a)** Workflow used to identify lncRNA-derived peptides enriched from cancer cells and tumors. We query the *de novo* sequencing-searched Class I-associated peptide pool against a database generated using six-frame translated lncRNAs compiled in LNCipedia database (right workflow). The statistical significance of our algorithm was determined by potential matching of the *de novo*-searched peptides against a “mock” database created by the randomly picked gene blocks (∼50,000 transcripts) from hg38, which resulted in an empirical p-value<1.0e^-5^ (left workflow). **b)** 44 lncRNA-derived peptides identified using our algorithm with their predicted HLA alleles and binding affinity. **c)** Log2 peptide intensities of database (DB) searched, *de novo* searched and lncRNA-derived peptides. **d)** The classification of source lncRNAs for the identified lncRNA-derived peptides into antisense, sense intronic and classic lncRNAs. **e)** LncRNA-derived peptides that match new open reading frame (ORF), introns of coding genes, and noncoding region. **f)** The top panel displayed a snapshot of IGV showing lncRNA PVT1-derived peptide FLLSSSLTL, identified in PC9, with chromosomal location alignment of RNA-seq of all 5 samples and peptide BLAT; the middle panel shows the ribo-seq searching results from GWIPS; the lower panel shows the predicted binding affinity of this peptide to HLA-A*02:06 that is expressed only in PC9 cells. **g and h)** Matched MS2 spectra of *in vivo* and synthetic lncRNA-derived peptides YSFPELTHL **(g)** and MEHVSPALP **(h)**, respectively. **i)** T2 cell-based HLA stability assay of three lncRNA peptides predicted to be HLA-A*02 binders, FLLSSSLTL, QEEAALKAL and SLHASLSTV, and the NY-ESO-1-dervied positive control peptide.

Next, we asked whether lncRNA-derived Class I-associated peptides had low abundance. We found that lncRNA-derived peptides were equally presented on Class I as all other DB and *de novo* sequencing-derived peptides. (**Figure 7c**). Based on LNCipedia classification, the source lncRNAs matched to 20 antisense genes, 12 sense intronic genes and 12 lncRNA genes (**Figure 7d**). Further BLAT analysis revealed 23 lncRNA coding regions had no overlap with any coding region on hg38, 5 lnRNAs matched to the introns of coding genes, and 16 matched to novel open reading frames (ORFs) due to the frameshift that intersected with exons of known protein coding genes, but with different start codons (**Figure 7e**). Frameshifted new ORFs have been suggested to be a rich source of neoantigens ^45^.

It remains unclear whether lncRNA can be translated to protein products (e.g., full length/truncated proteins, peptides), and importantly, presented to cell surface by HLA Class I. For in-depth illustration of our computational strategy, we analyzed further the lncRNA oncogene, *PVT1*-derived 9mer peptide, FLLSSSLTL, identified in PC9 cells. RNA-seq BAM files were converted to BED files for all five samples. The DNA coding sequence of the identified peptide, FLLSSSLTL, was retrieved from the *PVT1* nucleotide sequence in LNCipedia. The BLAT results of this 27-base pair sequence (i.e., TTC…CTT) was *in sillico* transcribed to RNA sequence (i.e., UUC…CUU) and further translated to peptide sequence FLLSSSLTL. Interestingly, we found that the source lncRNA were transcribed in all 5 samples, but a truncated version was transcribed in NCI-3795Mel. However, the peptide was only identified in PC9 cells, suggesting that translation of lncRNA-derived peptides may be cell and context-specific or the presentation of lnc-RNA-derived peptides by Class I is cell line or tumor-specific. We reasoned that the specificity to PC9 cells could be a result of the lack of specific HLA allele to present this A*02 restricted peptide (%Rank=0.11). Only PC9 and NCI-3795Mel express HLA-A*02 and NCI-3795Mel express a truncated version of FLLSSSLTL; this may explain why we only observed this peptide in PC9 cells. We further confirmed that the ORF containing this peptide coding sequence has been reported in the ribosome profiling data based on deep sequencing of ribosome protected mRNA fragments that can be visualized using GWIPS-viz (genome-wide information on protein synthesis)^46^ (**Figure 7f**). We further confirmed our MS identification of these lncRNA-derived peptides that are presented by MHC Class I. We used synthetic peptides and performed LC tandem MS to compare the MS2 spectra of synthetic peptides with those of the endogenous peptides. The MS2 spectra of lncRNA-derived endogenous peptides, YSFPELTHL derived from Lnc-ZNF804A, and MEHVSPALP, derived from Lnc-GALNT4, matched to the spectra of their synthetic counterparts (**Figure 7g-h**). Finally, we used the T2 cell-based HLA stability assay to confirm that 3 lncRNA-derived peptides were truly HLA-A*02 binders (**Figure 7i**). Taken together, we report a novel MS-based Class I-associated peptidome profiling platform for identification of lncRNA-derived peptides that are presented by HLA Class I.

## Discussion

Neoantigens are an attractive immunotherapeutic target because they specifically engage specific TCRs in T cells to eliminate tumor tissue while sparing nearby healthy tissues, unlike immune checkpoint blockades (e.g., PD1, PD-L1, CTLA-4) which non-specifically boost the immune response. Emerging evidence obtained from breast cancer ^47^, bladder cancer ^48^, melanoma ^49^ and lung cancer ^50^ studies suggests that cancer neoantigens may be ideal targets for ACT and therapeutic cancer vaccines. MS-based peptide-seq technology provides direct experimental evidence for a large number of HLA presented peptides. As such, this approach has become a robust and quick method of neoantigen discovery^14, 51, 52^. Nevertheless, a majority of studies have focused on “hot” tumors, such as melanoma or other high TMB lung cancer types^16, 53^. The “cold tumors” that have a paucity of TILs, tumors with low TMB such as EGFR mutant lung cancer and tumors with loss of neoantigen expression while on immunotherapy are relatively resistant to immune checkpoint therapy^54–57^. In this study, it was our intent to leverage proteogenomics and informatics to identify HLA Class I-presented peptide antigens, including neoantigens, common driver oncogenes, CG antigens, posttranslationally modified peptides and LncRNA - translated protein products, for immunotherapy in low TMB “cold” EGFR mutant lung adenocarcinoma. Moreover, we utilized targeted proteomics and HLA binding assays to further validate our proteogenomic findings.

To achieve this goal, we first verified the quality of our database-searched HLA Class I immuno-peptidome by confirming that a majority bind their corresponding HLA allele in the original sample. This is consistent with previous large-scale monoallelic HLA Class I epitope profiling studies^14, 58^. The NetMHCpan-predicted binding scores of the identified peptides favor 9 and 10mer peptides in comparison to longer peptides (**Figure 1f**). Our platform combined SNVs, INDELs and fusion variants to, for the first time identify 4 neoantigen-dervied peptides with a MAF less than 0.05, of which, peptide TEIGTIISV derived from PRRC2Cp.M2171I, has been associated with lung adenocarcinoma^59^ and reported as a lung cancer pathogenic variant in COSMIC. Since the germline data of the lung adenocarcinoma cell lines were not available, we identified several SNP-derived peptides. We acknowledge that truncal mutations in common oncogenes, such as *EGFR* and *KRAS*, derived Class I and Class II epitopes, may be the most attractive targets for ACT^8, 50^. We have reported the identification of somatic mutated peptides from the proteome of lung cancer patients, including a novel somatic mutated CDK12p.G879V peptide using similar methodology^60^. Although we identified 19 peptide epitopes derived from common oncogenes (**Figure 2g**), none of them contained somatic mutations.

Possible explanation of the identification of relatively few mutated neopeptides from known oncogene mutations include the absence of the cognate Class I allele and limitations of the data dependent acquisition (DDA) methodology for mass spectrometry-based sequencing of peptides when the mutated peptides are just a minute fraction of the total wild type peptides presented by HLA Class I. However, it is noted that the median MS-intensity of the few neoantigens we identified is very similar to that of wild type peptides (p-value>0.05) indicating that some neoantigens are robustly presented by Class I.

We have generated the most comprehensive CG antigen database to date by leveraging the human proteome atlas and multiple peer-reviewed CG antigen databases. This is a valuable resource which can be used to query CG antigens from other large cohort immunopeptidome studies. For instance, we identified LDHC (**Supplementary Figure 3a**) which, prior to this study, was almost exclusively observed in testis and an association with lung adenocarcinoma had only been suggested^61^. Given that CG antigens have been extensively investigated in melanoma^62, 63^, we have now revealed the CG antigen landscape in EGFR mutant lung cancer representing a “cold tumor”, unveiling novel Class I-presented peptides reported in our study. Interestingly, our results, contrary to previous observations^16^, show that CG antigen-derived peptide levels are not significantly correlated with mRNA and source protein expression (**Figure 4e**).

One of the advantages of a PEKAS database search is its pan-PTM search engine which unveils all PTMs in one search, requiring no prior knowledge of the potential types of modifications expected. To our knowledge, this study provides the deepest coverage of the PTM HLA peptidome, to date. Compared to previous profiling of PTM immunopeptides ^15, 16^, we, for the first time, systematically quantified the largest number of endogenous PTMs of HLA peptides in EGFR mutant lung adenocarcinoma. We demonstrated that unmodified peptides were significantly more abundant than their deaminated or methylated counterparts. However, this does not apply to the conversion of glutamate to pyroglutamate where a modification could lead to protein misfolding^33^. Generally, low peptide abundance or presentation in antigen presenting cells may hinder T cell recognition. It is possible that more abundant peptides are more likely represented as T cell epitopes. Although phosphorylation is the most dominant PTM known to modulate cellular function, we identified only a minor fraction of Class I-presented peptides phosphorylated; this could be due to the relatively large phosphate group which may not easily fit into the HLA binding groove. Researchers have reported only around two thousand unique phosphopeptides binding to 72 HLA alleles^64^. Also, phosphorylation, because it is a transient reversable PTM, generates less-stable peptides which are less likely good targets. This is in contrast to irreversible clinically relevant PTMs such as deamination which are more likely to generate neoepitopes. We discovered that a single peptide could have multiple PTMs concurrently (**Figure 3k**). Position +1 was the most frequently modified (**Figure 3d**), which implies that small modifications on a non-anchor position may not dramatically affect binding affinity. This is substantiated by comparing the predicted binding score of the modified only peptides with the modified/unmodified version. The modified only peptides are less subject to the NetMHCpan prediction, indicating these peptides may possess a unique HLA binding domain structure (**Figure 3i**) which compromises the prediction algorithm. Taken together, our results indicate the PTM peptides that are not predicted by any bioinformatic algorithm for HLA binding can be identified using mass spectrometry and these peptides may be a rich source as neo-epitopes.

Noncoding RNA is recognized as a rich resource for neoantigens ^65^, and MS-based proteogenomic platforms have been implemented to discover non-canonical nonpeptides^17^. Laumont and colleagues suggested that noncoding regions are the main source of neoantigens ^41^. In contrast to previous studies where MS spectra were mapped *in silico* to all potential reading frames of FASTA derived RNA-seq data, we leveraged deep learning methodology of *de novo* sequencing of peptides in PEAKS studio to query against the largest annotated LncRNA database (LNCipedia.org). Our approach extends the previous studies in many ways. Conventionally, an RNA-seq translated FASTA database contains an extremely large number of protein variant entries because a majority of the human genome is non-coding; as a result, a significant number of false positive matched peptides is expected. However, in this study, we utilized a relatively small LNCipedia database to match the high quality *de novo* searched peptides which have been stringently filtered by peptide length and HLA binding prediction scores. In addition to the average local confidence (ALC) score estimation by the PEAKS *de novo* search, we validated the significance of our findings by generating an empirical p-value when compared to a “mock” gene dataset. We also selectively validated the presence of the source lncRNAs in the ribosome profiling data searching the GWIPS database, suggesting that these lncRNAs are indeed translated on the ribosome machinery. To further ensure that the identified LncRNA-derived peptides were indeed not a part of a known expressed protein, we manually checked and visualized that the source lncRNA of each of these novel peptides was indeed present in the total RNAseq data from the same cell line and tumor and did not overlap with any coding region or was not due to a frameshift. Synthetic peptide validation and T2 cell HLA binding assays further confirmed that the identified LncRNA-derived peptides were indeed presented and had high affinity to the HLA proteins. Taken together, the pipeline we developed in this study could be readily applied to any type of cancer to identify lncRNA-derived peptides presented by HLA Class I.

Deep learning *de novo* search engines imbedded in PEAKS provided high accuracy to detect peptides that normally are missed in a database search. Our pipeline evaluates in-depth the *de novo* only peptides in the immunopeptidome context. The validity of this approach for HLA peptidome profiling and neoantigen identification is underscored by several findings in this study. The binding motif of the 9mer *de novo*-only peptides shared almost 100% similarity to the DB searched peptides **(Figure 6g-6j**). Half of the neoantigens identified in the proteogenomic pipeline were also retrieved in the *de novo*-only peptides using normal human proteome database (**Figure 6k**). Single neopeptides retrieved from the *de novo* search, proteogenomic DB search and synthetic standard were a perfect match to their MS/MS sequencing spectra (**Figure 6l**).

Overall, we report the largest characterization of potential cancer epitopes in EGFR-mutant lung adenocarcinoma to date. The combination of genomics, proteomics and cutting-edge informatics allows us to develop this in-depth immunopeptidome-based cancer epitope profiling pipeline. Our results prove that low TMB tumors possess as many potential immunotherapy targetable epitopes as “hot” or high TMB tumors. We provide a valuable resource of mutant EGFR lung cancer specific epitopes for the design of targeted immunotherapy and cancer vaccines.

## Methods

### Subjects and cell lines

The primary lung tumor was obtained at rapid autopsy from left lower lobe lung of osimertinib treated patient RA007, a 70-year old male, with a primary EGFR^L858R^ SNV. The patient has been consented with Institutional Review Board (IRB) approved protocol 13-C-0131 (NCG01851395) entitled “A Pilot Study of Inpatient Hospice with Procurement of Tissue on Expiration in Thoracic Malignancies.” The clinical trial which patient RA007 was enrolled was approved by the NCI IRB with the local protocol number 13C0131. The patient was offered hospice treatment with life expectancy less than 3 months at CCR, NCI. The rapid autopsies were initiated within 3 hours upon patient death. The clinical trial details are available under <u>NCG02759835</u> and have been thoroughly described by our group previously ^66^.

The two primary melanoma cell lines, NCI-3784Mel and NCI-3795Mel were obtained from Surgery Branch, National Cancer Institute. The clinical trial identifier of these primary cells is NCT00068003 “Harvesting Cells for Experimental Cancer Treatments”, and NCI-784Mel has been reported previously^37^. The melanoma cells were cultured in high glucose Dulbecco’s modified eagle medium supplemented with 20% fetal bovine serum (FBS). Mutant EGFR lung adenocarcinoma cell lines, PC9 and H1975, were purchased from ATCC. The lung adenocarcinoma cells were maintained in RPMI-1640 medium supplemented with 10% FBS. The tumor was immediately snap frozen in liquid nitrogen upon lesion excision. The T2 (174 x CEM.T2) cell line, purchased from ATCC, was maintained in ATCC-formulated Iscove’s modified Dulbecco’s medium supplemented with 10% FBS.

### HLA peptide enrichment and purification

For melanoma and lung adenocarcinoma cell line HLA peptide enrichment, 2.0 × 10^8^ cells were harvested in 4ml ice-cold lysis buffer (20mM Tris-HCl pH=8.5, 100mM NaCl, 1 mM EDTA. 1% triton X-100 supplemented with Halt 1:100 protease Inhibitor cocktail Cat. No 78430, Thermo Scientific). After 30min on ice, lysates were subjected to needle sonication for 30s. Approximately 30mg snap-frozen lung tumor tissue was homogenized in 4ml ice-cold lysis buffer for 30s at 4°C using the Qiagen TissueLyser II. Cell/tissue lysates were centrifuged for 2 hours at 4°C at 20,000g, and the supernatant used in subsequent experiments. HLA-peptides complexes were isolated by interacting with 0.5mg HLA Class I pan antibody clone W6/32 (BioXcell, West Lebanon, NH) pre-coupled to 200μl slurry of protein A/G PLUS agarose resin (Santa Cruz Biotechnology) overnight at 4°C with constant rotation. Agarose beads were then washed three times with ice-cold lysis buffer (without triton and protease inhibitors), followed by two washes in ice-cold 20mM Tris-HCl (pH=8.5), then one wash in ice-cold HPLC grade water. Complexes were eluted four times with 0.15% trifluoroacetic acid (TFA) in water at room temperature and combined. To purify immunopeptides, HLA-peptide complexes were loaded on preconditioned 50mg C18 desalting columns (Sigma Millipore), then followed three 0.1% TFA in water washes. HLA peptides were then eluted with 40% acetonitrile (ACN) in 0.1% TFA. Purified peptides were lyophilized at -80°C for 2hours, then, reconstituted in 0.1%TFA, 2%ACN loading buffer for MS analysis.

### Tandem mass spectrometry-based peptide sequencing and validation

Estimated 1μg purified and desalted HLA peptides were loaded to a 2cm nano Acclaim trap column (Cat. No 164535), then were separated on a 25cm EASY-spray reverse phase column (Cat. No ES802A) for 90min effective gradient with 4-35% 0.1% formic acid in ACN on an Ultimate 3000 Nano liquid chromatography instrument (Thermo Scientific, Waltham, MA). The separated peptides flew to an Orbitrap Q-Exactive HF mass spectrometer (Thermo Scientific, Waltham, MA) with discovery mode for data acquisition. The MS1 full scan (375-1650 m/z) was set to 120,000 resolution, and top 15 most abundant peptides per cycle were subsequently fragmentated by high collision dissociation (HCD). To identify only non-tryptic digested neutral HLA Class I-associated peptides, typical with 8-14 amino acid residues, only charge state 1-4 was included for MS2 scans. The MS2 sequencing scans acquired the peptide fragments at 30,000 resolution and 200ms max injection time window. The dynamic exclusion was set at 20s.

Selected nonpeptides and LncRNA-derived peptides were *in vitro* synthesized by GenScript (Piscataway, NJ). Selected melanoma CG antigen-derived peptides were synthesized by the Peptide Synthesis and Antigen Discovery Core, Surgery Branch, NCI. Synthetic peptides were serial diluted to 1pmol/μl in 0.1%TFA in water, then, all peptides pooled and subjected to MS analysis. To maintain retention time matching, the same LC-MS/MS settings were used for the spectra validation analysis and the discovery MS.

### Database and *de novo* search of MS raw files

Patient and cell line specific protein sequence databases were first generated. The aligned whole exome sequencing BAM files from patient blood (germline) or tumor tissue/cell lines were used to retrieve the variants call format (VCF) files using HaplotypeCaller ^67^, and the intermediate VCF files were further annotated by SnpEff ^68^which filtered out only nonsynonymous on exome regions including SNVs and Indels. Similarly, BBduk was used to remove adapter sequences and low-quality reads from paired-end FASTQ files, which were then used as input for STAR-Fusion ^69^. The final VCF files were *in silico* translated to sample specific protein sequence libraries (FASTA files) using QUILTS ^70^, which were merged with refseq hg38 converted human proteome database.

The database search of MS raw files was carried out by PEAKS studio ^39^ (Bioinformatics Solutions) using the patient/cell line specific databases described above. In the PEAKS searching engine, no enzyme digestion was selected because HLA peptides are natural peptides without artificial digestion. Importantly, the unique PEAKS built-in functions, pan-PTMs including 650 different variable modifications and *de novo* searching were used. The precursor mass tolerance was set to 15 ppm and fragment ion tolerance was set to 0.5 Da. For the mutant neoepitope screening, the false discovery rate (FDR), estimated by decoy-fusion database, was chosen at 0.05. For the wt peptides screening (e.g., CG antigen, LncRNA and PTMs), FDR was set to 0.01. For the *de novo* search, we applied very stringent criteria, a) the average local confidence (ALC) score (substituted for FDR estimation) of each peptides must be >50%; b) the lowest %rank, predicted by NetMHCpan4.0, of each peptides against their corresponding HLA molecules in the same sample must be <2.0. The detected raw peptide intensity was log2 transformed for further statistical analysis.

### HLA Class I typing

Seq2HLA package ^21^ was used to call the four-digit HLA Class I typing from the WES of all samples. Additionally, HLA typing (six-digit) of the patient donors of the two primary melanoma cell lines were performed by the Department of Transfusion Medicine (DTM) of Clinical Center at National Institutes of Health. Results obtained by WES-based HLA typing and the patient HLA testing at DTM/CCR/NIH were consistent.

### Whole-exome and total RNA sequencing and total proteome profiling

WES and RNA-seq was performed as described previously ^71^. Briefly, the genomic DNA and total RNA of the cell lines and tumor were extracted and sent to the NGS core facility at National Cancer Institute (NCI) Frederick National Laboratory. The samples were sequenced as 2 × 126 nt paired end reads with Illumina HiSeq2500 sequencers with >100million reads per sample. The raw FASTQ files were aligned to hg38 by TopHat (v2.0.13) ^72^, the alignment BAM files were used for downstream variant calling. For the samples with a corresponding germline specimen, Strelka (v1.0.10) was used for somatic variant calling, and “pass” entries were assigned as somatic variants ^73^. The total RNA seq of H1975 (n=3) were normalized by Reads Per Kilobase of transcript, per Million mapped reads (RPKM) and quantified.

Roughly 0.2mg of the cell lysate of H1975 were reduced and alkylated, and further digested by MS grade trypsin/lysC at 37°C for 16 hours. The tryptic peptides were fractionated by an off-line high-pH reverse phase liquid chromatography (LC) into 12 fractions, each of which was subjected to a 120 min gradient separation on a nanoLC and analyzed by an Orbitrap Q-Exactive HF using discovery mode. The 12 MS raw files were searched by MaxQuant (v1.5.7.4) against a Uniprot Human database.

### Generation of Cancer Germline (CG) antigen database

We compiled a CG antigens database using existing antigens reported in CTDatabase (http://www.cta.lncc.br/) and Cancer Antigenic Peptide Database (https://caped.icp.ucl.ac.be/), and literature searches. Lastly, we manually verified the tissue expression of each CG antigen gene in testis and other tissues in the Human Protein Atlas (https://www.proteinatlas.org/) to ensure their RNA and protein expression are mainly in testis but not in other tissues.

### Immunoblotting of HLA Class I antigens

One million cells from each cell line were lysed in ice-cold modified RIPA buffer for 30 min. The cell lysate was spun down at 20,000 g for 15 min at 4°C, and supernatant was transferred whose protein concentration was determined by the bicinchoninic acid protein assay. Aliquoted 10μg protein from each cell line was load to SDS-PAGE. Subsequently, separated proteins were transferred from gel to polyvinylidene fluoride membrane and incubated with primary anti-HLA Class I mouse HRP mono-antibody, at 1;5000, (EMR8-5, Funakoshi) overnight at 4C, then briefly incubated with SuperSignal HRP substrates (Thermo Scientific) before imaging. The membranes were exposed for 5 seconds, and images were acquired by Odyssey Fc imager (LI-COR Biosciences).

### Flow cytometry analysis for HLA Class I expression

One million cells were collected in 100μl FACs buffer (PBS+5%FBS). After 30min blocking, half of the cells were incubated with FITC anti-HLA Class I, W6/32 (Cat. No 311404, BioLegend) for 30min at 4°C, and washed twice with FACs buffer. Flow cytometry was carried out on CytoFLEX platform (Beckman), and 20,000 events were collected per sample. The post-analyses and statistics were conducted by FlowJo (v10.6.2).

### HLA binding affinity T2 cell assay

The TAP-deficient HLA-A2 only expression T2 cells (unable to process endogenous HLA ligands) were counted and 1.0 million cells/ml were suspended in growth medium, then plated in 6-well tissue culture plates (2ml/well). The testing synthetic peptides and NY-ESO-1 peptide (positive control) were reconstituted in DMSO, and further diluted to a final concentration of 10μM with growth medium then incubated with T2 cells at 37°C with 5% CO2 for 12 hours. Each testing peptide and positive control were performed in triplicate and negative control (DMSO) in duplicate. After 24 hours, flow cytometry was performed as described above to detect the cell surface total HLA Class I expression.

### Identification of HLA presented LncRNA-derived peptides

For the genomic data processing, quality control was done on RNA-seq FASTQ files by first assessing read quality using FASTQC (v0.11.8), then by removing adapter sequences and low-quality reads using BBduk, part of the BBTools package (v38.42). The resulting FASTQ files were aligned against the RefSeq human genome version hg38 using STAR (v1.3.4) ^74^, then sorted using the Samtools mappings sorter (v 1.1.1) ^75^. Duplicate reads were then removed using Picard MarkDuplicates (v2.1.1). These alignments, sorting and de-duplication steps were run on the DNANexus platform. The resulting BAM file was converted to BED format using the bamtobed tool from the BEDTools suite (v2.29.0) ^76^. The BEDTools intersect tool was used to find the intersection between the tumor or cell-line BED file and the LNCipedia (www.lncipedia.org) hg38 high-confidence (HC) BED file (v5.2) which contains ∼50,000 annotated lncRNAs ^44^. The result was a BED file of the matched expressed genes in the sample transcriptome.

For the proteogenomic pipeline development, the *de novo* searched 8-14mer peptides were queried against a 6-frame translated FASTA file of the all the lncRNAs on LNCipedia (v5.2). The unique IDs of the lncRNAs that have a peptide matched to were used to retrieve the lncRNA entries in the LNCipedia HC BED file (RefSeq hg38). The chromosome notation in the peptide-matched lncRNA BED file and the bed file of the sample transcriptome were made to have the same chromosome notation (e.g. “CM00663.2” into “chr1”). Then, BEDTools intersect was used to find the intersection of the peptide-matched lncRNAs and the LNCipedia -matched expressed genes in the sample transcriptome BED file in order to find which genes in the tumor or cell line transcriptome matched certain query peptides. The resulting intersection bed file was inspected for overlaps with the original cell line or tumor transcriptome and the human genome (RefSeq hg38) in Integrative Genomics Viewer (2.5.3) ^77^. Any peptides whose corresponding matched lncRNA also matched to the noncoding region or intron region but not the coding exon region of the human genome were kept as potential LncRNA-derived peptides.

For the ribosome sequencing, we utilized a well-recognized ribo-seq genome browser (https://gwips.ucc.ie/index.html) which compiles 46 ribosome profiles ^46^. The lncRNA containing ORFs were manually searched on the GWIPS to confirm the lncRNA-derived peptide coding regions RNAs were bound to ribosome.

For the empirical p-value evaluation, we randomly sampled 50,000 non-overlapping genomic sequences, each with length 2000 base pair (the mean length of a lncRNA from the LNCipeida datase) uniformly from the human reference genome sequence to be used as mock lncRNAs. Each of these mock lncRNAs gave rise to 6 reading frames, from which, at most 600 potential peptides were extracted. Comparing this mock peptide set to the MS data through our computational pipeline resulted in 65 matches. In comparison, LNCipedia database includes 107039 lncRNA transcripts from ∼50,000 distinct lncRNAs, where each lncRNA may give rise to multiple transcripts primarily due to alternative splicing. From this larger set of lncRNA transcripts (and using all 6 reading frames) the extracted potential peptides resulted in 195 matches from the mass spec data. For a pessimistic (larger than the correct value) estimate on the p-value for this outcome, we set the probability of a chance match of a potential peptide to p = 66/(50000*12000/avg.pept.length). The probability that there are exactly k matches in the entire set of potential peptides from the real lncRNA transcripts (there are n = 107000*12000/avg.pept.length such potential peptides) based on the null assumption is: Then the empirical p-value can be calculated:

## Data availability

The raw mass spectrometry-based sequencing files (.raw) of HLA Class I immunopeptides profiling of the NCI-3784mel (n=5), NCI-3795mel (n=5), PC9 (n=6), H1975 (n=3) and NCI-RA007 (n=5) have been deposited in the online repository of proteomics data, ProteomeXchange (http://proteomecentral.proteomexchange.org/cgi/GetDataset), with the an accession number PXD019774. The raw files can be reviewed during the peer-review process using a reviewer account (Username: and Password: weujUujk). The patient/sample specific fasta files have been deposited on Synapse.org with a Synapse ID: syn22227675 and the public data is accessed at https://www.synapse.org/#!Synapse:syn22227675/.

## Code availability

The original Python scripts of the identification of HLA presented LncRNA-derived peptides are public available on the Github repositories at: https://github.com/YueAndyQi/lncRNA_immunopeptidome_Scripts. The bioinformatics analysis of study was performed on the Biowulf Linux cluster at the NIH (https://hpc.nih.gov/docs/userguide.html).

## Acknowledgments

We thank Drs. Paul F. Robbins and Maria R. Parkhurst (Surgery Branch, CCR, NCI, NIH) for their generous assistance in melanoma CG peptides synthesis and helpful discussions in melanoma CG peptide identification. We also thank Dr. Jared Gartner (Surgery Branch, CCR, NCI, NIH) for providing assistance in analysis of RNA-sequencing datasets of the melanoma cell lines.

## Funding source

This study was supported by the Intramural Research Program of Center of Cancer Research (CCR), National Cancer Institute (NCI) of the U.S. National Institutes of Health.

## Author contributions

Y.A.Q., and U.G. concepted the research proposal and designed the study. Y.A.Q., T.K.M., K.D.N. and C.A. performed the cell culture and sample preparation for proteomic analysis. C.A., and K.H., and J.C.Y. generated the melanoma primary cells, performed the RNA-seq and called somatic mutations of the melanoma cells. Y.A.Q., and X. Z. conducted the mass spectrometry analysis and database search. D.M., and J.K. constructed the cancer germline antigen database. Y.A.Q., V.M., and S.G. performed the next generation sequencing data analysis and HLA typing. Y.A.Q., C.A., K.H., and J.C.Y. discussed and performed T2 cell-based HLA stability assay. Y.A.Q., V.M., M.H.E., and C.S. generated the computational platform of LncRNA-derived peptides identification. Y.A.Q., drafted the manuscript. Y.A.Q., C.M.C., and U.G. edited and revised the manuscript. Y.A.Q., and U.G. supervised the overall experimental and data analysis of this project. All authors discussed, reviewed, and approved the manuscript.

**Supplementary Table 1:** Summary of database and *de novo* searched HLA-Class I associated peptides.**Table 1:** HLA tying of cancer cell lines and lung adenocarcinoma tumor.

**Supplementary Table 1:** Summary of database and *de novo* searched HLA-Class I associated peptides.

**Supplementary Table 2:** Somatic mutation lists of two melanoma primary cell lines and one lung adenocarcinoma tumor tissue.

**Supplementary Table 3:** Identified high-confidence oncogene/tumor suppressor-derived peptides.

**Supplementary Table 4:** Summary of pan-PTM searched HLA Class I associated peptides.

**Supplementary Table 5:** In-house generated cancer germline antigen database. **Supplementary Table 6:** Summary of identified cancer germline antigen-derived peptides.

**Supplementary Table 7:** Summary of identified long noncoding RNA-derived peptides.

## Competing interests

U.G. has a clinical trial agreement (CTA) with AstraZeneca and had received research funding from AstraZeneca and Aurigene. U.G. is currently an employee of Bristol Myers Squibb. The other authors have no conflicts of interest to report.

---

**Supplementary Figure 1.**
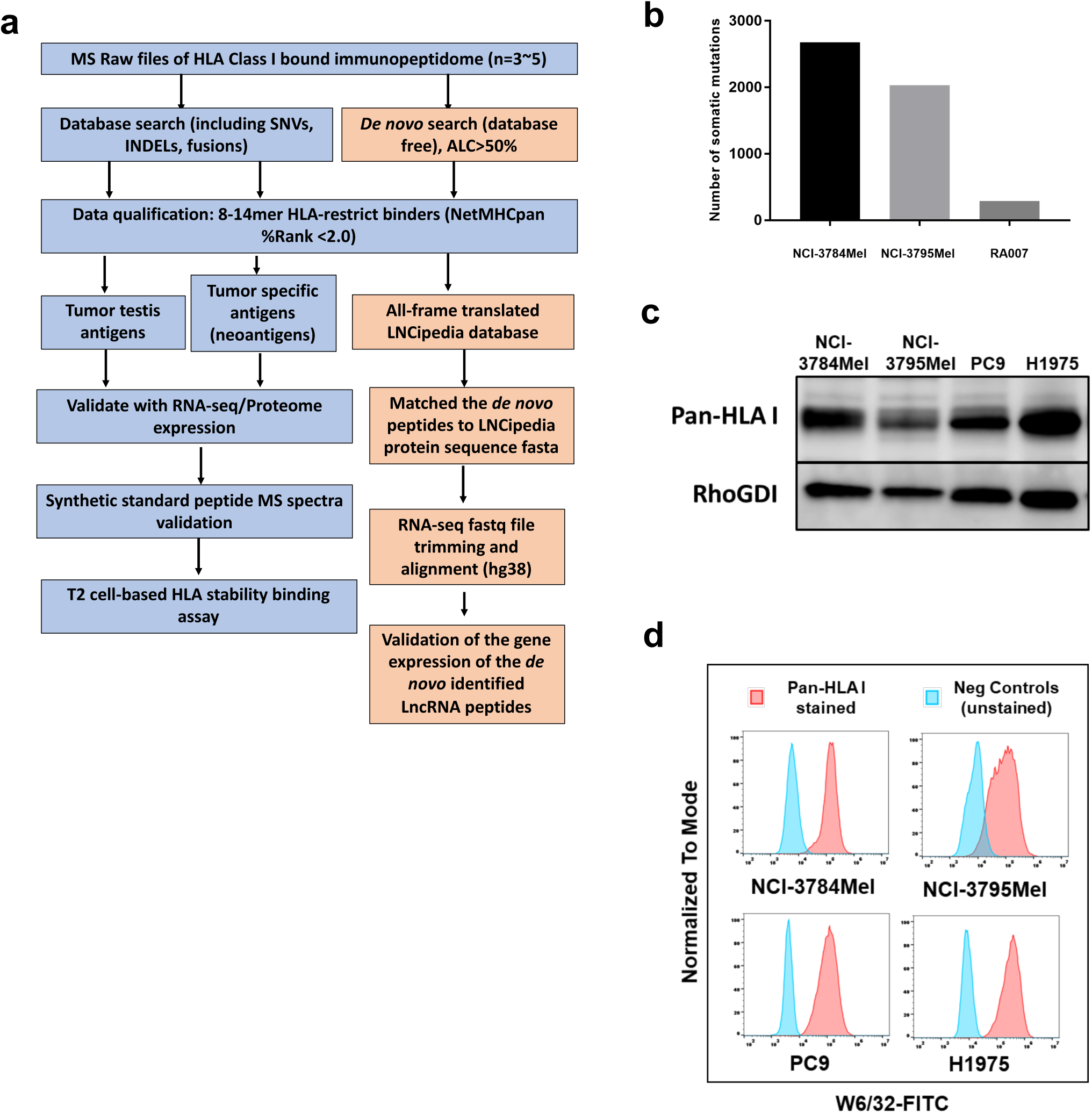
Detailed strategic plan of current proteogenomic analysis and immunoblotting and FACs of HLA molecules. **a)** Detailed strategic workflow of MS raw file processing, database search, cancer antigen-derived peptides identification and validation. **b)** Number of somatic mutations identified from NCI patient melanoma tumor derived cell lines (NCI-3784Mel and NCI-3795Mel) and an EGFR mutant lung adenocarcinoma tumor, procured at autopsy, from a patient treated with osimertinib (NCI-RA007). **c)** Immunoblots of total HLA Class I protein expression using pan HLA Class I (antibody clone W/32). **d)** Flow cytometry analysis of cell surface HLA Class I protein expression.

**Supplementary Figure 2.**
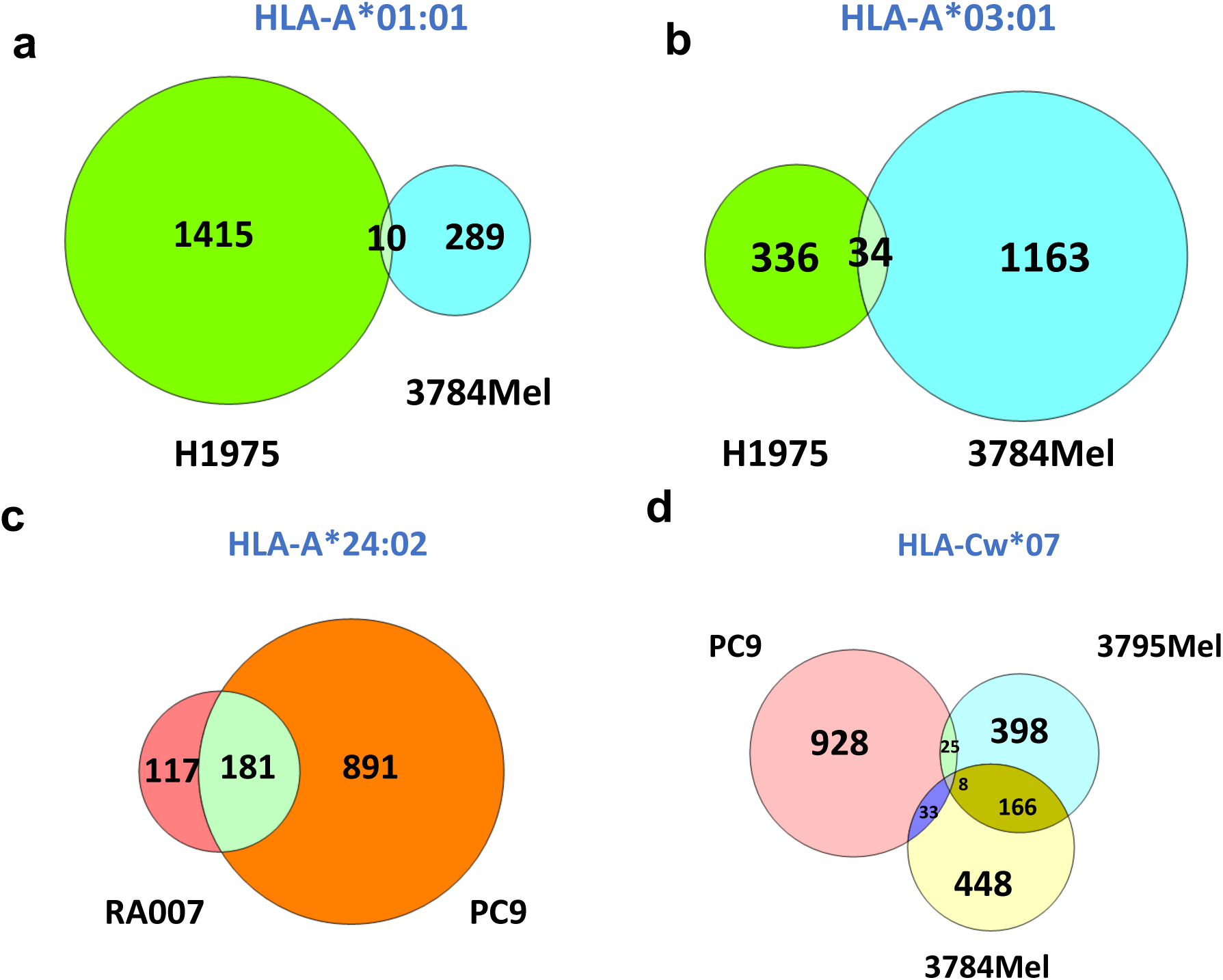
HLA restrictive binding peptidome varies in different cancers. **a-b)** Venn diagram shows the intersects of predicted HLA-A*01:01 **(a)** or HLA-A*03:01 **(b)** binding immunopeptides identified in H1975 and NCI-3784Mel, respectively. **c)** Venn diagram shows the intersects of predicted HLA-A*24:02 binding immunopeptides identified in NCI-RA007 and PC9. **d)** Venn diagram shows the intersects of predicted HLA-C*07 binding immunopeptides identified in PC9, NCI-3784Mel and NCI-3795Mel.

**Supplementary Figure 3.**
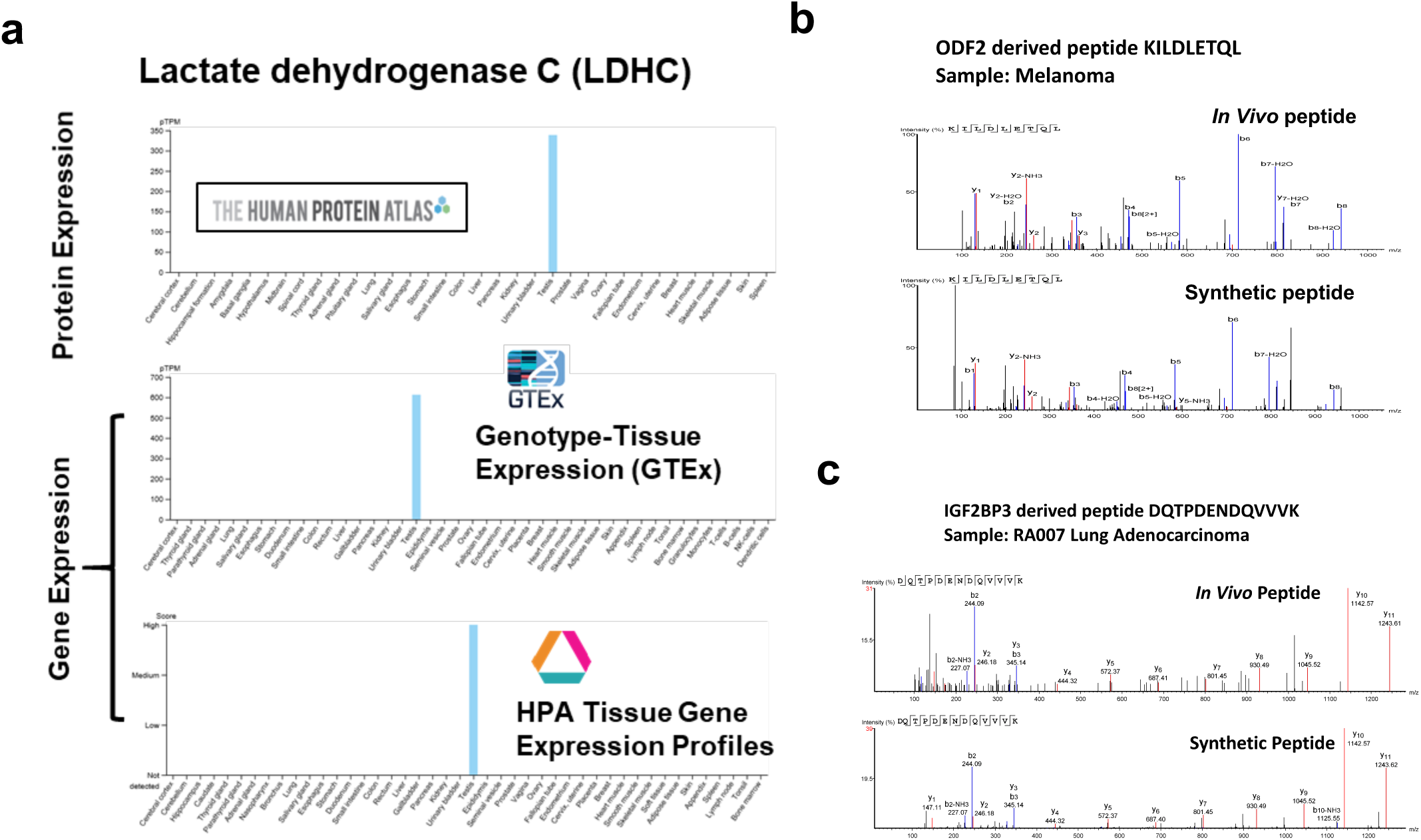
Customized CG antigen database and synthetic peptide validation. **a)** Representative example of a CG antigen-lactate dehydrogenase C (LDHC) showing expression, at the RNA and protein levels, in testis but not in other organs (data from The Human Proteome Atlas). **b)** Matched MS2 spectra of endogenous outer dense fiber of sperm tails 2 (ODF2) derived peptide KILDLETQL identified in NCI-3784Mel and its synthesized counterpart. **c)** Matched MS2 spectra of endogenous Insulin-like growth factor 2 mRNA-binding protein 3 (IGF2BP3)-derived peptide DQTPDENDQVVVK identified in NCI-RA007 and its synthesized counterpart.

**Supplementary Figure 4.**
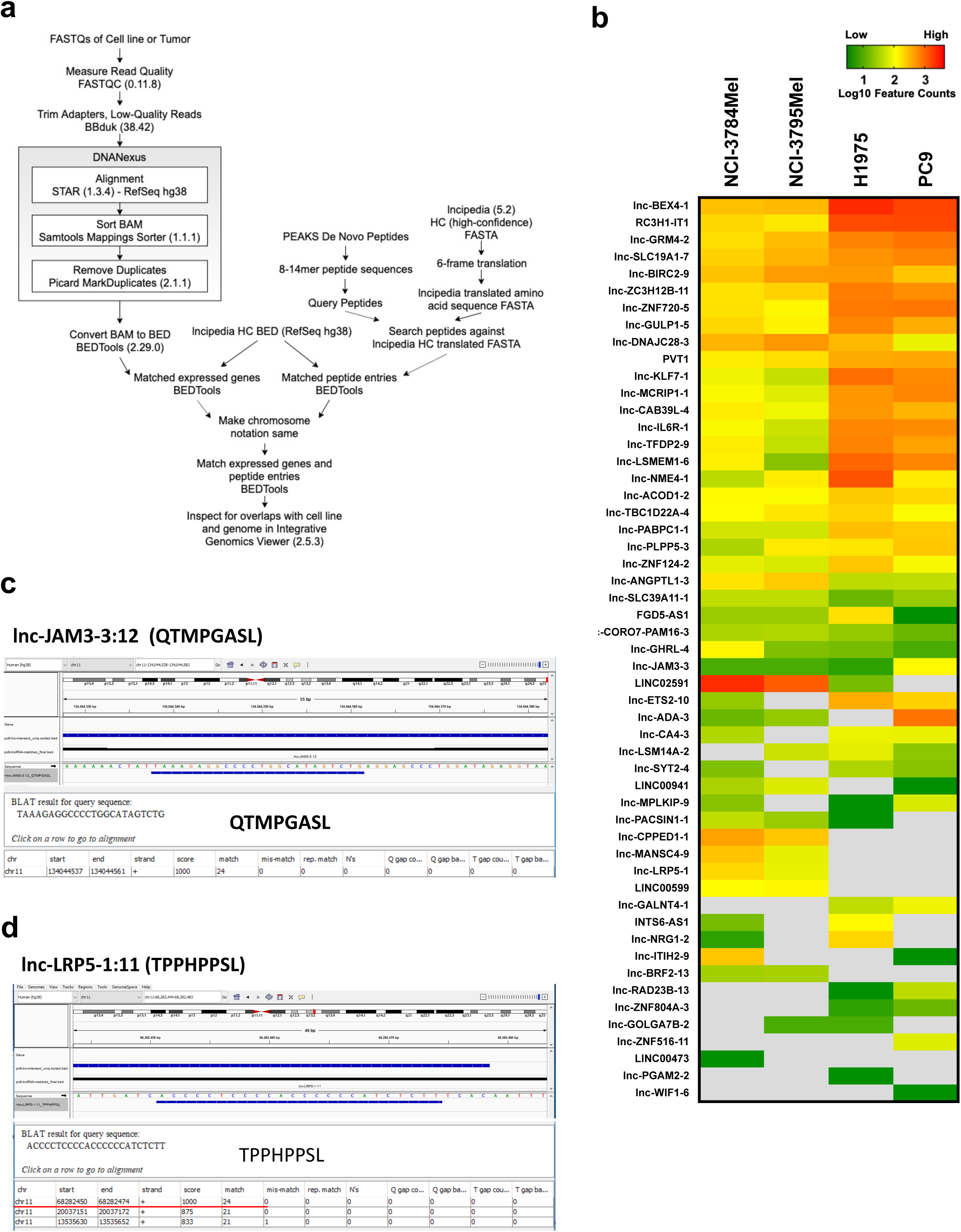
Detailed LncRNA peptide identification computational workflow and manual inspection and visualization of discovered LncRNA peptides. **a)** Detailed bioinformatic workflow of lncRNA peptides identification. **b**) Heatmap of source lncRNA gene expression in log10 feature counts. **c-d)** IGV snapshot showing Lnc-JAM3-3:12-derived peptide QTMPAGSL **(c)** and Lnc-LRP5-1:11-derived peptide TPPHPPSL **(d)**, identified in PC9, with chromosomal location alignment of hg38 and RNA-seq. The peptide BLAT results were shown in the bottom panel.

## Notes

https://figshare.com/articles/journal_contribution/Proteogenomic_analysis_unveils_the_HLA_Class_I_presented_immunopeptidome_in_melanoma_and_EGFR_mutant_lung_adenocarcinoma/12759977

### Competing Interest Statement

U.G is a current employee of Bristol Myers Squibb. U.G. has a Clinical Trial Agreement with AstraZeneca and received research funding from AstraZeneca and Aurigene.

